# Targeting Aurora kinases as essential cell cycle regulators to deliver multi-stage antimalarials against *Plasmodium falciparum*

**DOI:** 10.1101/2025.07.10.663886

**Authors:** Henrico Langeveld, Keletso Maepa, Marché Maree, Jessica L. Thibaud, Nicolaas Salomane, Rosie Bridgwater, Mufuliat T. Famodimu, Luiz C. Godoy, Charisse Flerida A. Pasaje, Nonlawat Boonyalai, Mariana Laureano de Souza, Justin Fong, Tayla Rabie, Rensu P. Theart, Sonja Ghidelli-Disse, Jacquin C. Niles, Marcus C. S. Lee, Elizabeth A. Winzeler, Michael J. Delves, Kelly Chibale, Kathryn J. Wicht, Lauren B. Coulson, Lyn-Marié Birkholtz

**Author notes:** Corresponding author: Lyn-Marié Birkholtz.

## Abstract

Kinases that play critical roles in the development and adaptation of *Plasmodium falciparum* present novel opportunities for chemotherapeutic intervention. Of particular interest are mitotic kinases that regulate the proliferation of the parasites by controlling nuclear division, segregation and cytokinesis. We evaluated the potential of human Aurora kinase (Aur) inhibitors to inhibit *P. falciparum* development by targeting members of the Aurora-related kinase (Ark) family in this parasite. Several human AurB inhibitors exhibited multistage potency (<250 nM) against all proliferative stages of parasite development, including asexual blood stages, liver schizonts and male gametes. Among the most potent compounds, hesperadin and AT83 exhibit >1000x selectivity towards the parasite without concerns about mammalian cell toxicity. Importantly, we identified *Pf*Ark1 as the principal vulnerable Ark family member, with specific inhibition of *Pf*Ark1 as the primary target for hesperadin and the human anaplastic lymphoma kinase (ALK) inhibitor TAE684. Hesperadin’s whole-cell and protein activity validates it as a unique *Pf*Ark1 tool compound. Inhibition of *Pf*Ark1 results in the parasite’s inability to complete mitotic processes, presenting with unsegregated, multi-lobed nuclei caused by aberrant microtubule organization. This suggests that *Pf*Ark1 is the main Aur mitotic kinase in proliferative stages of *Plasmodium*, characterized by bifunctional AurA and B activity. This paves the way for drug discovery campaigns based on hesperadin targeting *Pf*Ark1.

## Introduction

*Plasmodia spp.* parasites demonstrate pathogenic success due to their complex life cycle, alternating between non-proliferative stages, where the cell cycle is quiescent, and stages characterized by rapid cell division events that lead to massive parasite population expansion^1,2^. *Plasmodium falciparum,* the causative agent of the most severe form of malaria^3^, undergoes three unique and specialized cell division events. During hepatic and intra-erythrocytic schizogony within the human host, a haploid parasite undergoes multiple rounds of closed asynchronous mitosis and karyokinesis, followed by a singular and synchronized cytokinesis event to produce a segmented schizonts^4,5^. An equally unique cell division event occurs within the mosquito host, where male gametocytes (1n) undergo three rounds of rapid DNA replication (exflagellation) to generate eight flagellated male gametes (8n) in just ∼15 min^6,7^. The parasite’s ability to undergo rapid asexual replication is a key factor in its pathogenic success but requires extraordinary control. A detailed mechanistic understanding of the role players in regulating the parasite’s atypical cell cycle in *Plasmodium* could lead to novel antimalarial therapeutic agents.

Cell cycle machinery and regulators, such as protein kinases (PKs), are crucial for accurate progression through various checkpoints in mammalian cells. Several conserved Ser/Thr mitotic PKs are considered primary regulators of the mitotic process, including the ‘Never In Mitosis’ kinases (NIMA/Neks), Polo-like kinases, and Aurora kinases (Aur). The Aur family is highly conserved amongst eukaryotes, with members identified, amongst others, in yeasts (Ipl1), humans (*Hs*AurA, B, and C), *Toxoplasma gondii* (*Tg*Ark1–3)^8^, *Trypanosoma brucei* (*Tb*AUK1)^9^. In *P. falciparum,* three aurora-related kinases exist (*Pf*Ark1–3)^10,11^. Aurora kinases contribute to the assembly and disassembly of mitotic and meiotic centrosomes, regulating spindle pole structure and dynamics, chromosome segregation, and cellular fission during cytokinesis. While Aur members are differentiated functionally depending on their localization, delocalization can cause moonlighting effects between *Hs*AurA and B^12^, although direct compensation for the loss of activity is not evident^13^. During mitosis, *Hs*AurA (‘polar’ Aur) localizes to the centrosome and spindle poles, and upon binding of microtubule-associated protein TPX2, regulates centrosome maturation, separation, and microtubule spindle formation. *Hs*AurB as ‘equatorial’ Aur (with INCENP, survivin, and borealin), forms the chromosome passenger complex (CPC) as master controller of cell division, localized to centromeres, kinetochores, and the spindle midzone, allowing AurB to govern chromosome condensation, kinetochore attachment, sister chromatid segregation, and cytokinesis.

*Pf*Arks are implicated in critically regulating cell cycle progression of *Plasmodium* based on 1) the essentiality of all three *Pf*Arks to asexual proliferation (schizogony)^14^, 2) the unique expression patterns of the *Pf*Arks during cell cycle arrest and re-entry^15^ and 3) distinct, highly specific, and exclusive spatiotemporal associations during ABS schizogony^16^. Although *P. falciparum* lacks a canonical centrosome, it possesses a microtubule-organizing centre (MTOC) characterized by a centriolar plaque (CP) embedded in the nuclear envelope, with inner CP (intranuclear body) and outer CP domains (cytoplasmic body)^17^. The CPs harbor several validated centrosomal proteins, including centrin and γ-tubulin, and facilitate microtubule (MT) nucleation. *Pf*Ark1 (PF3D7_0605300) and *Pf*Ark2 (PF3D7_0309200) are associated with MTOCs during schizogony^16^, with *Pf*Ark1 localizing to the outer CP domains of duplicated MTOCs in nuclei primed for division (similar to a ‘polar’ Aur^10^) while *Pf*Ark2 is additionally proposed to localize to kinetochores, akin to an ‘equatorial’ Aur^16^. *Pf*Ark3 (PF3D7_1356800) is found only in segmented nuclei associated with subpellicular microtubules (SPMTs) as cytosolic microtubules in merozoites, suggesting a role in cytokinesis^16^.

The deregulation of *Hs*Aurs (especially *Hs*AurA and *Hs*AurB) has been linked to cancer and tumorigenesis, making them attractive targets for anticancer therapeutic strategies^18^. Several inhibitors selectively target either *Hs*AurA or *Hs*AurB or have dual or pan-reactive abilities. *Hs*AurA inhibition leads to defects in mitotic spindle assembly and ultimately causes spindle checkpoint-dependent mitotic arrest, cell cycle exit and apoptosis^19^. On the other hand, *Hs*AurB inhibition causes abnormal chromosome alignment and overrides the mitotic spindle checkpoint, causing polyploidy, failure of cytokinesis, and endoreduplication^20^.

Although several kinase families have been validated as antimalarial targets (e.g. *Pf*PI4K^21^, *Pf*PKG^22^ and *Pf*CLK3^23^), the unique requirement of Ark members for parasite proliferation processes has not been extensively explored to identify novel inhibitors specifically targeting this kinase family. Previous studies have shown that the *Hs*AurB-specific inhibitor hesperadin exhibits potent *in vitro* activity against *P. falciparum, T. brucei* and *Leishmania donovani*^24–26^. We aslo demonstrated that hesperadin treatment leads to mutations in *Pf*Ark1 that confer resistance^27^. However, evidence of direct target engagement and inhibition of the *Pf*Ark members is lacking.

Here, we provide an in-depth evaluation of *Pf*Ark inhibition and its influence on parasite survival. We systematically assessed *Hs*Aur inhibitors targeting all Aur classes to identify compounds that could be repurposed as antimalarials. We identify Ark inhibitors with multistage activity against proliferative stages of the parasite, including asexual blood stage (ABS) parasites, male gametes and liver schizonts, correlating with the required function of Arks in cell proliferation events. Biochemical characterization showed the specific and sensitive inhibition of *Pf*Ark1 as the primary target of the inhibitors, with exquisite potency seen for hesperadin and NVP-TAE684 (TAE684). Hesperadin is a *bona fide* inhibitor of only *Pf*Ark1, without targeting other kinases or exhibiting additional pleiotropic activity associated with the inhibition of hemozoin formation (the crystalline byproduct of hemoglobin digestion), as is the case with TAE684 and the other kinase inhibitors studied. A unique aspartate residue in the *Pf*Ark1 active site confers selectivity to hesperadin inhibition over the mammalian Aur and *Pf*Ark2, which have a lysine residue at the equivalent position. *Pf*Ark1 is shown to be the main Ark family member involved in mitotic processes, and interference with its activity results in aberrant nuclear division with no clear microtubule nucleation at the CPs in both ABS parasites and male gametes. These findings provide chemical validation of *Pf*Ark1 as a druggable target, with hesperadin presenting a starting point for further drug discovery initiatives.

## Results

### Aur inhibitors demonstrate effectiveness against the replicative stages of *P. falciparum* parasites

A set of commercially available compounds was chosen based on their specificity and potency to the *Hs*Aur members. The inhibitors were first assessed against various life cycle stages of *Plasmodium* parasites *in vitro* to determine their multistage anti*plasmodium* activity (Figure 1, Supplementary Figure S1). Less than half of the *Hs*AurA inhibitors targeted ABS proliferation of drug-sensitive NF54 *P. falciparum* parasites with IC_50_ values <5 μM (Figure 1A, Supplementary file S1). By contrast, both *Hs*AurB, most of the dual-active inhibitors and five of the six pan-active Aur inhibitors showed activity at this concentration. Six compounds (AAi I & Aki III [targeting *Hs*AurA^28,29^], hesperadin & AZD-1152 [targeting *Hs*AurB^30,31^], AT83 & ZM-39 [dual *Hs*AurA&B^32,33^]) were all potent at <1 μM. All inhibitors that exhibit specificity towards *Hs*AurB, even when developed as dual *Hs*AurA&B, are generally more potent against *P. falciparum* ABS, with an IC_50_ of 1.5 nM for hesperadin (Supplementary Figure S2A), and <250 nM for AT83 and ZM-39. The eight most active compounds retained activity against multidrug-resistant *Pf*Dd2 and *Pf*K1 strains with ≤2-fold variance in IC_50_ from *Pf*NF54 (Supplementary Figure S2B). TAE684 (Novartis’ first-generation anaplastic lymphoma kinase (ALK)-specific inhibitor) was also included in this assay as it was proposed to target a *Pf*Ark based on Kinobead competitive pulldown data^34^ (Supplementary Figure S2C), with TAE684 also presenting potent activity against ABS parasites at IC_50_ = 280 ± 13 nM.

**Figure 1:**
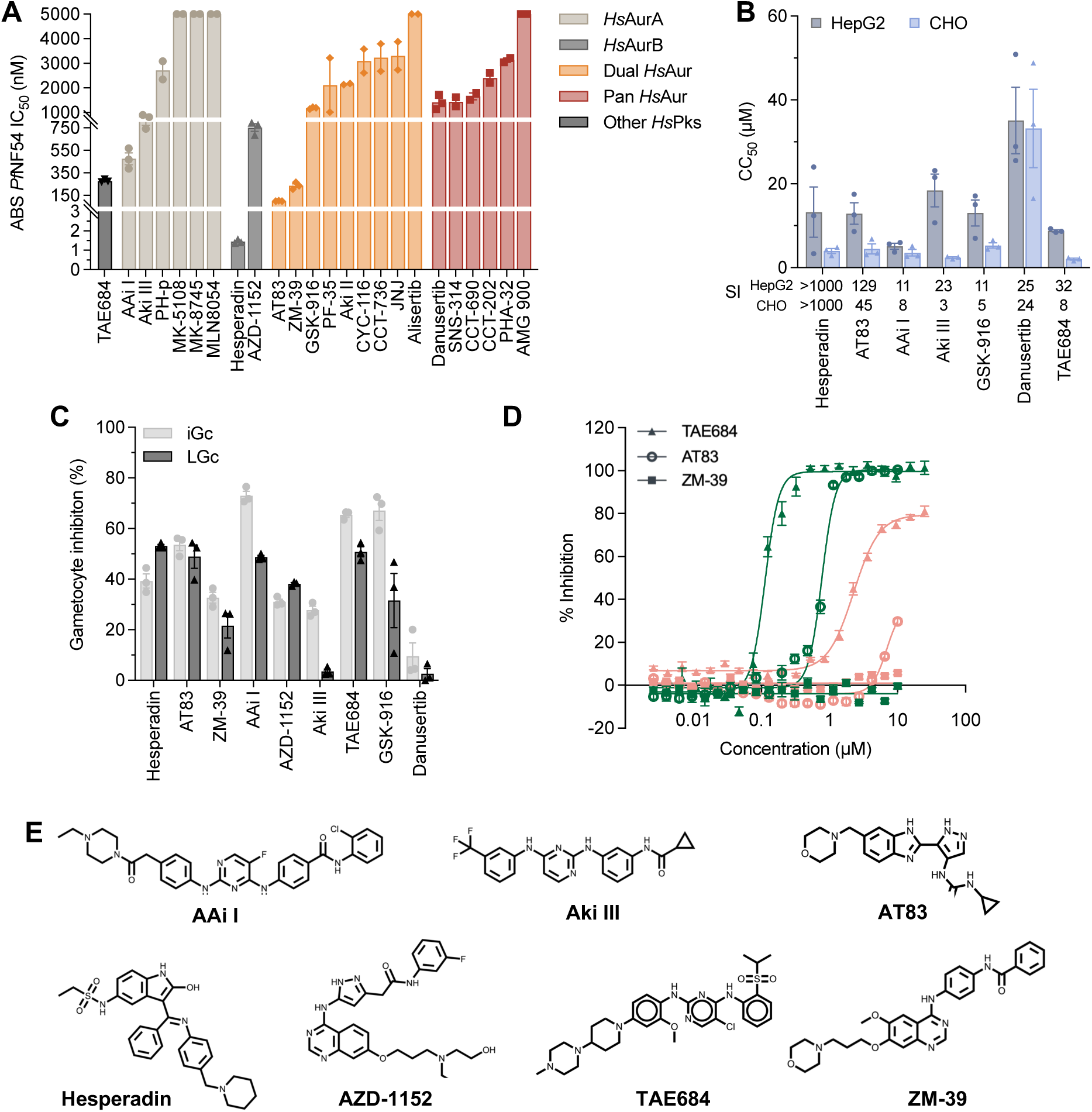
Activity profile of Aurora kinase class anticancer inhibitors across the life cycle of *P. falciparum* parasites. **A)** Activity (IC_50_) of inhibitors (grouped based on specificity against human Aurora kinases) against *P. falciparum* drug-sensitive (*Pf*NF54) asexual intra-erythrocytic parasites. Values obtained for the most active inhibitors are from three independent biological replicates, each performed in technical triplicate (n=3, mean ± S.E.), while the remainder is from two independent biological replicates, each performed in technical triplicate (n = 2, mean ± S.E.). **B)** Cytotoxicity (CC_50_) of selected inhibitors against HepG2 and CHO cell lines. The selectivity Index (SI = HepG2 CC_50_/*Pf*NF54 IC_50_ and CHO CC_50_/*Pf*NF54 IC_50_) is indicated below the graph (n=3, mean ± S.E.). C) Single point activity profile of selected inhibitors at 5 µM against immature (iGc, II-III) and late-stage (LGc, IV/V) gametocytes determined by measuring luminescence of recombinant *P. falciparum* parasites expressing luciferase (n=3, mean ± S.E.). **D)** Activity (IC₅₀) against male gamete formation (green) and female gametes (pink) for TAE684, AT83 and ZM-39 (n=3, mean ± S.E.). **E)** Structures of the seven most active compounds selected.

With selectivity and mammalian cell toxicity often being concerns with kinase inhibitors, we evaluated the activity of the active compounds against hepatocellular carcinoma (HepG2) and Chinese Hamster Ovary (CHO) cell lines (Figure 1B). Several compounds exhibited cytotoxicity with selectivity indices (SI) <10, including GSK-916, AAi I, Aki III and TAE684. The pan-active inhibitor danusertib demonstrated the most pronounced cytotoxic effect against both lines. However, *Hs*AurB inhibitors hesperadin and AZD-1152 showed distinct selectivity in targeting *P. falciparum* compared to mammalian cell lines, indicating differentiation in action in the parasite, with SI >1000 and >100-fold, respectively. Similarly, the dual *Hs*AurA&B inhibitors, AT83 and ZM-39, also showed SI >100-fold.

We subsequently evaluated the ability of the compounds active on NF54 ABS parasites to target additional life cycle stages of *P. falciparum* parasites. All selected inhibitors displayed minimal (<50% inhibition at 5 µM) gametocytocidal activity, with only AAi I, GSK-916 and TAE684 showing ∼70% inhibition of immature gametocyte viability (>80% stage II/III) whilst unable to kill mature gametocytes effectively (Figure 1C). This finding was not unexpected, as gametocytes are non-proliferative cells, although all three *Pf*Arks are expressed during gametocytogenesis on a transcript and protein level^16,35^. However, against male gamete formation, a process that similarly requires DNA replication and cell division, hesperadin (IC_50_ of 10 nM^36^), AT83 (786 ± 16 nM) and TAE684 (116 ± 17 nM) prevented male gamete exflagellation, whereas only TAE684 had any appreciable activity against female gametes (IC_50_ = 2.3 ± 0.05 μM), with the rest inactive (>10 µM) (Figure 2D). Confirming the preference for proliferative forms, hesperadin and TAE684 display a low nanomolar (IC_50_ <200 nM) potency against *P. berghei* liver schizonts (Supplementary Figure S2D). Taken together, these data reveal that a focused set of *Hs*Aur-specific inhibitors demonstrates potent and selective anti*plasmodium* activity, preferentially targeting proliferative parasite stages related to cell cycle division, while exhibiting an acceptable margin of cytotoxicity.

**Figure 2:**
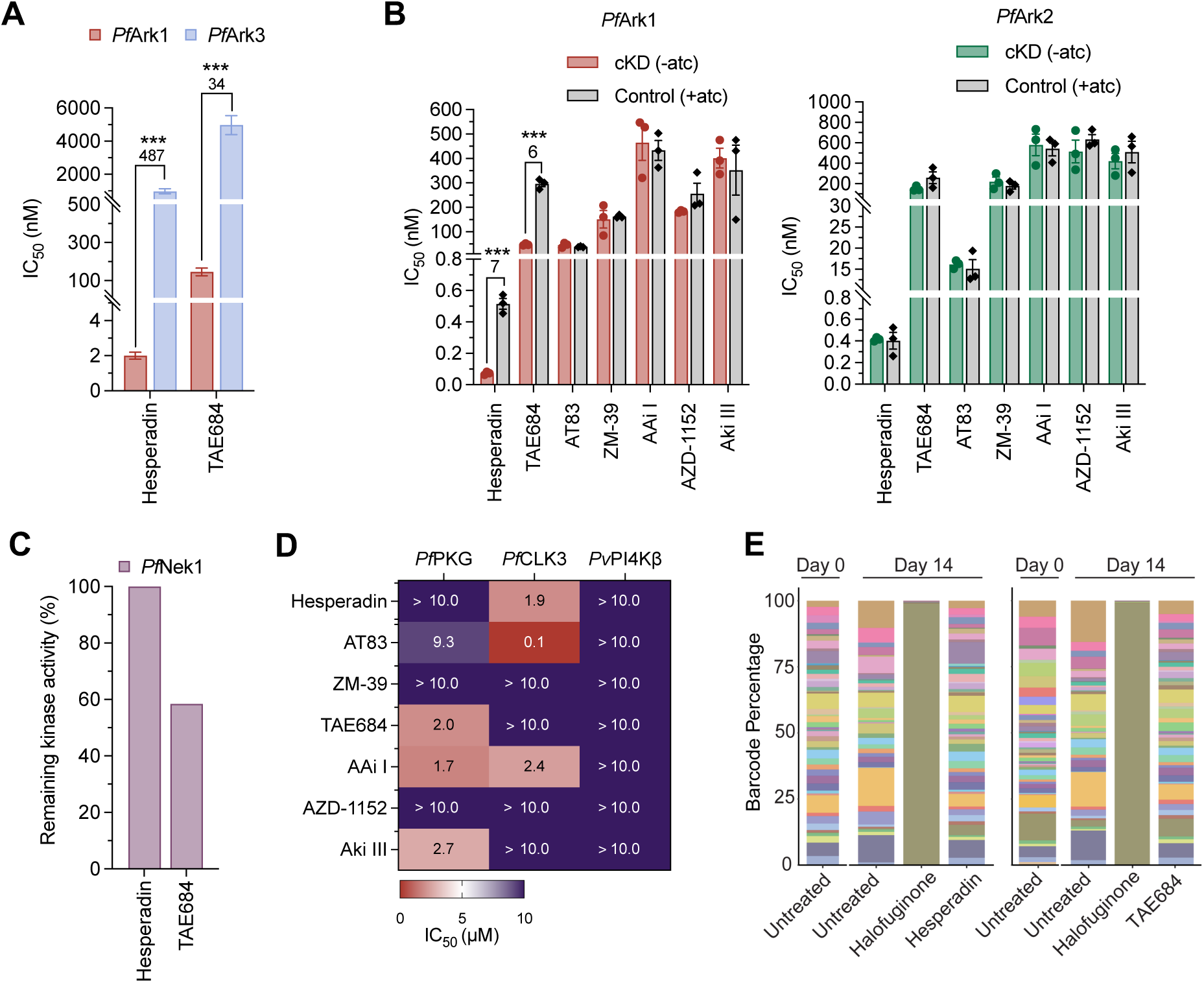
Identification and validation of *Plasmodium* Aurora-related kinase as the primary target. **A)** Activity (IC_50_) against recombinant *Pf*Ark1 and *Pf*Ark3 proteins, using a three-hybrid split-luciferase competitive binding assay (KinaseSeeker™). **B)** Effect of conditional knockdown (cKD) of *Pf*Ark1 and *Pf*Ark2 on parasite sensitivity relative to control conditions in the presence of high aTc. Representative dose-response curves are presented for each cKD parasite line (n=3, mean ± S.E.) with an unpaired two-tailed t-test, \**p* < 0.05; \*\**p* < 0.01; \*\*\**p* < 0.001. **C)** Single-point activity profile of selected inhibitors at 1 µM against recombinant *Pf*Nek1 (KinaseSeeker™). D) Inhibitory activity against recombinant *Pf*PKG, *Pf*CLK3, and *Pv*PI4Kβ. The mean IC_50_ values ± SD were calculated using two independent experiments (n=2), each with technical duplicates using the ADP-Glo Kinase Assay. **E)** Stacked bar plots illustrating barcode populations on days 0 and 14 for no drug, drug control (halofuginone), and selected inhibitors.

### *Pf*Ark1 was identified and validated as a novel *Plasmodium* kinase target

Given that the selected inhibitors were developed and optimized to specifically target different *Hs*Aur mitotic kinases, we aimed to correlate the whole-cell inhibition of the proliferative stages of *P. falciparum* to the inhibition of *Pf*Ark proteins (Figure 2). We first used the KinaseSeeker™ competitive binding assay to determine the inhibitory activity against *Pf*Ark1 and *Pf*Ark3^37^. Among the selected inhibitors tested, AT83 and AAi I had a marginal effect (AT83: 30% inhibition of *Pf*Ark1, AAi I: 40% inhibition of *Pf*Ark3, Supplementary Figure S3A). However, hesperadin and TAE684 exhibited potent activity against *Pf*Ark1, with IC₅₀ values of 2 ± 0.2 and 146 ± 20 nM, respectively (Figure 2A). Notably, both hesperadin and TAE684 had a significant preference towards *Pf*Ark1, with hesperadin ∼480-fold more active against *Pf*Ark1 than *Pf*Ark3 (*p*=0.0004 and *p=*0.0001, respectively, n=3, unpaired Student’s t-test).

We evaluated the involvement of *Pf*Ark2 inhibition by determining the loss of activity of the compounds against conditional knockdown (cKD) lines of *P. falciparum* for either *Pf*Ark1 or *Pf*Ark2^38^. Most compounds did not show a change in activity against the cKD of either *Pf*Ark1 or *Pf*Ark2 (Figure 2B). However, cKD of *Pf*Ark1 resulted in an increased sensitivity to hesperadin and TAE684, as evidenced by a significant >5-fold decrease in the IC_50_ values compared to the wild-type control (*p=*0.0002 *and p=*0.00003, respectively, n=3, unpaired Student’s t-test; Figure 2B, Supplementary Figure S3B). Conversely, there was no change in the IC_50_ value for these compounds in the *Pf*Ark2 cKD. This further supports *Pf*Ark1 as the primary protein target for hesperadin and TAE684 (Figure 2C).

The specificity towards *Pf*Ark1 was confirmed by evaluating the ability of the compounds to inhibit other Ser/Thr or lipid kinases. This included inhibition of *Pf*Nek1, as hesperadin resistance selections previously yielded mutations in this gene, suggesting an epistatic interaction between *Pf*Ark1 and *Pf*Nek1^27^, and *P. berghei* Nek1 exhibits similar spatiotemporal associations with the outer CP domain as *Pf*Ark1^38^. However, hesperadin did not inhibit *Pf*Nek1 activity, even at 1 μM, with only TAE684 showing a marginal 40% effect on this protein (Figure 2C). The inhibitors did not exhibit noteworthy activity against three other validated kinase antimalarial targets (*Pf*PKG, *Pf*CLK3, or *Pv*PI4Kβ) (Figure 2D), except for AT83, which inhibited *Pf*CLK3 with an IC_50_ of 100 nM. Additionally, hesperadin, TAE684 and AT83 were evaluated for their ability to inhibit the proliferation of a set of resistant *P. falciparum* parasites (including resistance mutants for *Pf*PI4k and *Pf*CLK3) using the antimalarial resistome barcode sequencing (AReBar) assay^39^. All three compounds killed all of the resistant lines in the platform (Figure 2E, Supplementary Figure S3C and Supplementary File S2), indicating no cross-resistance with known antimalarial resistance mechanisms and a novel mode of action.

These data indicate that the specific inhibition of *Pf*Ark1 within the family of Arks in *P. falciparum* by hesperadin and TAE684 isa the primary driver of ABS anti*plasmodium* activity. This correlates with the specific increase in abundance of this protein (over *Pf*Ark2) peaking at ∼ 28–34 hpi^40^ in preparation for its availability during schizogony (Supplementary Figure S3D).

### The activity of the additional *Hs*Aur inhibitors is not associated with Ark inhibition

Since all the compounds selected for this study were based on their activity against *Hs*Aur, but we could only convincingly show that hesperadin and TAE684 target *Pf*Arks, we explored alternative mechanisms of activity for some of the most potent inhibitors (AT83, ZM-39, Aki I, AAi III and AZD-1152). Several kinase inhibitors inhibit hemozoin formation due to structural similarities in the presence of multiple heteroaromatic rings, planar structures, and basic centers^41^. Evaluation of the ability to block formation of synthetic β-hematin (βH) *in vitro* in a cell-free detergent-mediated Nonidet P-40 (NP-40) assay^42^ indicated that AAi I, Aki III, AZD-1152 and ZM-39 (but not AT83) were potent inhibitors of βH formation (IC_50_ <20 μM), similar to the positive control chloroquine (CQ) (Figure 3A). This could implicate inhibition of hemozoin formation as a primary mode of action for these compounds. Interestingly, TAE684 displayed some effect against βH formation (IC_50_ of 41.4 ± 2.3 µM, Figure 3A), whereas hesperadin was ∼3-fold less active (IC_50_ of 126.3 ± 42.0 µM). The inhibition of βH formation translated to inhibition of intracellular hemozoin formation for both ZM-39 and TAE684^43^, which caused a significant increase in free heme (*p=*0.00000002 and *p=*0.0003, respectively, n=3, unpaired Student’s t-test), accompanied by a simultaneous decrease in hemozoin formation (*p=*0.00001 and *p=*0.00000003, respectively, n=3, unpaired Student’s t-test) (Figure 3B). This effect was not observed in the hesperadin treatment, with no change in either heme or hemozoin levels, implying that hesperadin does not affect hemozoin formation in the parasite (Supplementary Figure S4A).

**Figure 3:**
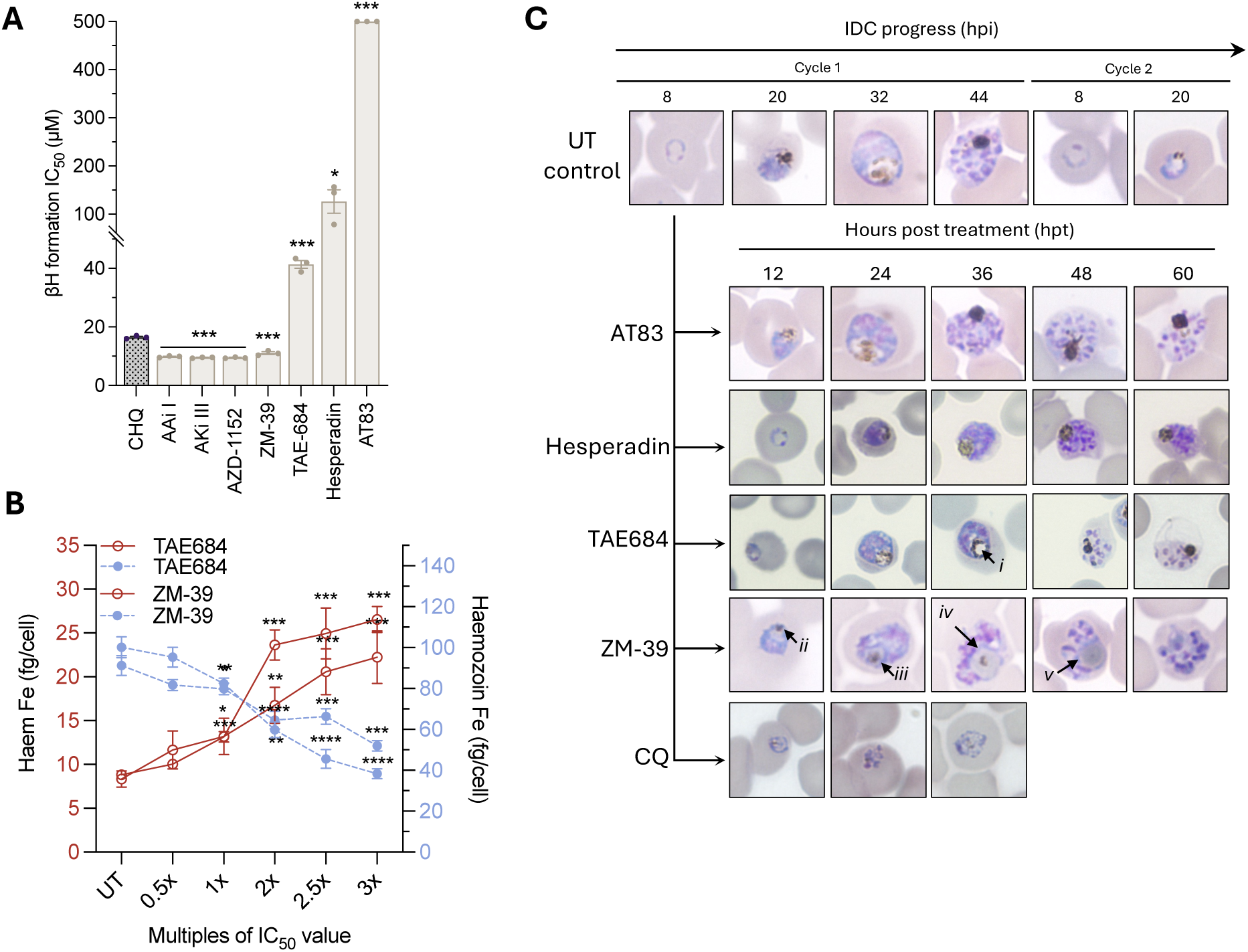
Phenotypic effect of potent *Hs*Aur inhibitors on intra-erythrocytic development. **A)** Measuring the ability of the inhibitors to interfere with the formation of synthetic hemozoin, βH, *in vitro* in a cell-free detergent-mediated NP-40 assay. Error bars represent ± SD for technical triplicates with an unpaired two-tailed Student’s *t*-test. Exact *p*-values provided in \**p* < 0.05; \*\**p* < 0.01, \*\*\**p* < 0.001, \*\*\*\**p* < 0.0001. **B)** Dose-dependent changes in the heme Fe levels from intracellularly-extracted fractions of hemozoin under ZM-39 and TAE684 treatment. **C)** Phenotypic response of ABS *Pf*NF54 parasites exposed to AT83 (3xIC_50_, ∼600 nM), ZM-39 (3xIC_50_, ∼750 nM), TAE684 (3xIC_50_, ∼900 nM) and hesperadin (IC_99_, ∼3 µM). Parasite morphology was observed at 12 h intervals using thin blood smears and indicated enlarged food vacuoles (*i*, *iv* and *v*), and small hemozoin crystals (*ii* and *iii*).

This was confirmed by a distinct phenotypic morphology, where a decreased hemozoin crystal size (Figure 3C, *i* & *ii*) and an enlarged food vacuole-like structure (*iii* & i*v*) were observed for ZM-39 treatment within the first 24 h post-treatment (hpt). Importantly, the TAE684 treatment had a distinctly different phenotype, whereas hemozoin formation persisted for the first 24 h, but thereafter, a distinct vacuolar structure was present, and aberrant schizonts were formed. The morphological abnormalities associated with the food vacuole were not present in either hesperadin-or AT83-treated parasites. For both situations, parasites normally progressed and entered schizogony, however, schizonts were abnormal, particularly following hesperadin treatment, and persisted for 60 hpt, indicating an arrested state and functional impairment in the completion of schizogony (Figure 3C, Supplementary Figure S4B). Co-treatment of TAE684 or ZM-39 with chloroquine, a known hemozoin formation inhibitor, showed an additive or indifferent effect, as evaluated by fixed-ratio isobologram analysis, with ΣFIC_50_ values of 1.3 and 1.4, respectively. By contrast, hesperadin was antagonistic to chloroquine (ΣFIC_50_ 1.6) as well as to TAE684 (ΣFIC_50_ 3.5) (Supplementary Figure S5). Taken together, the data suggest that TAE684 exhibits polypharmacology, involving both the inhibition of hemozoin formation and *Pf*Ark1 inhibition, but importantly, hesperadin has *Pf*Ark1 as its singular target.

### Structural understanding of inhibitor-target interactions in the ATP-binding site of *Pf*Ark1

The specificity of hesperadin, an ATP-competitive inhibitor, against *Pf*Ark1 was mechanistically investigated. *Pf*Ark1 shares ∼34% identity with mammalian AurA and B, with both the ATP-binding signature and S/T protein kinase sites well conserved, including the catalytic lysine residue, Lys61 (Figure 4A, Supplementary Figure S5A). However, *Pf*Ark1 exhibits several critical changes in both the ATP binding site and the active site relative to both mammalian and protozoan Aur proteins, including alterations of two conserved Lys residues, one to Asp (at position 40) and Ala (at position 42), something only seen for the *T. gondii* Ark1, but not for *Pf*Ark2 (Supplementary Figure S5B). Additionally, there are changes in the gatekeeper residue from Leu to a bulkier and more flexible Met at position 109 and an Ala to Cys change at position 112 within the hinge region (Figure 4A).

**Figure 4:**
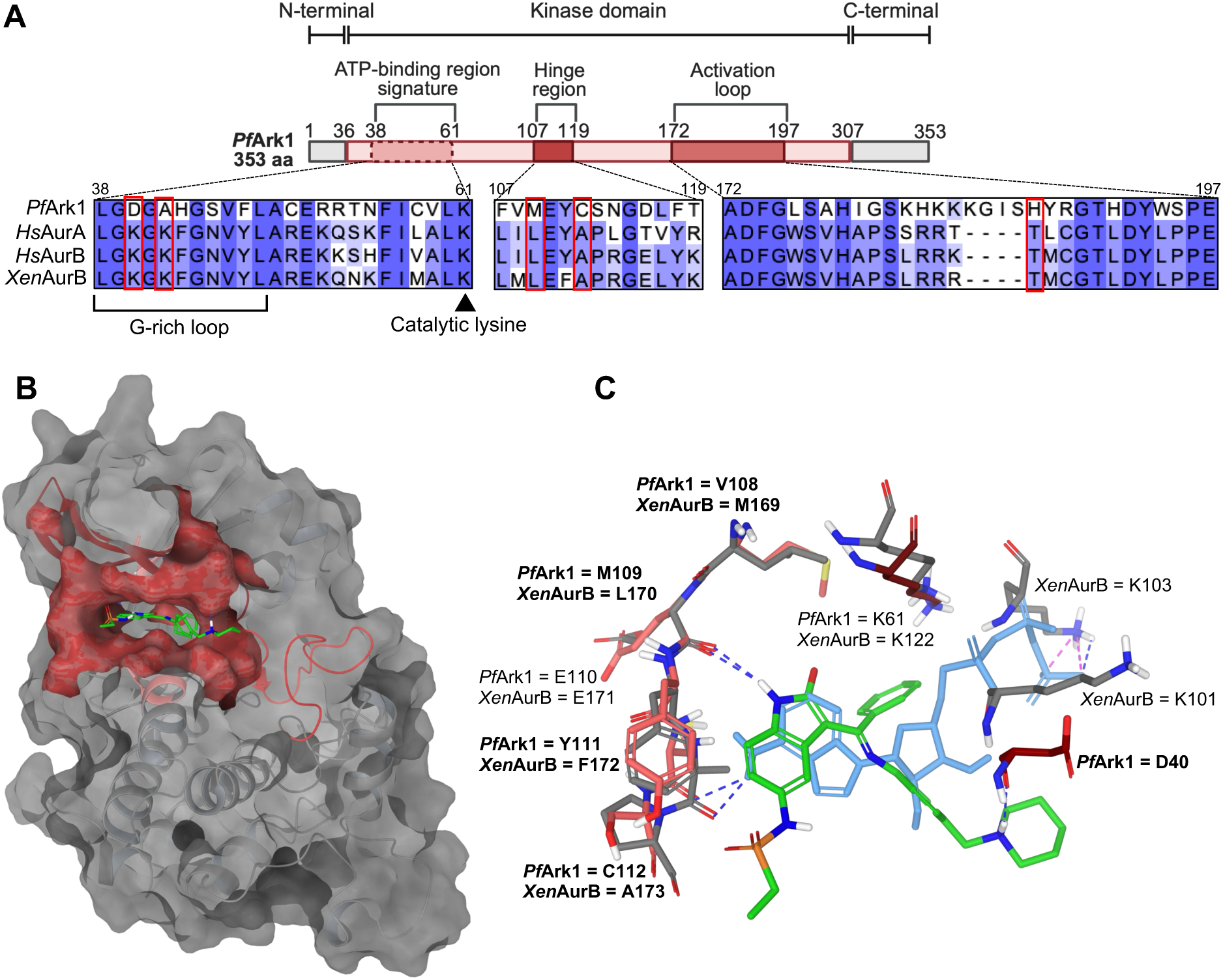
*In silico* modelling predicts protein-inhibitor interactions in the active site of *Pf*Ark-1. **A)** Diagram of key characteristics of *Pf*Ark1 protein along with a protein sequence alignment of mammalian and *P. falciparum* aurora kinases. **(B and C)** Hesperadin binding pose in *Pf*Ark1 (green), overlayed with the binding pose of ATP (light blue), and the *Xenopus laevis* AurB crystal structure (2BFY, grey), with residue differences indicated in bold. Selected main chains and side chains of key residues conserved within the kinase domain are displayed (red and pink are for *Pf*Ark1, while grey is for *Xen*AurB) with hydrogen bonds displayed as blue and salt bridge bonds as pink dashed lines.

*In silico* molecular docking studies revealed that hesperadin binds within the ATP-binding pocket of *Pf*Ark1 but in a different pose to that observed for *Hs*AurB and in the *Xenopus laevis* AurB co-crystallized with hesperadin^44^ (Figure 4B and C). Hesperadin’s indolinone moiety and sulphonamide group form hydrogen bonds, respectively, with the key conserved hinge region residues Glu110 and Tyr111. This causes the central phenyl to point into the active pocket of *Pf*Ark1, which would displace the α-phosphate of ATP and prevent its interaction with the catalytic lysine (Lys61) (Figure 4C). Indeed, analogues lacking the bulky phenyl are not as active as hesperadin^24^. Hesperadin is additionally stabilized in the ATP binding site pocket by an H-bond between its piperidine ring and the unique Asp40 found in the *Plasmodium* enzyme (Figure 4C). *Pf*Ark1 is differentiated from *Pf*Ark2 in this position, with the latter containing a more conserved K-N modification compared to the *Hs*Aur, which does not accommodate stabilization of hesperadin to the same extent as in *Pf*Ark1. This data provides clarity on the selectivity of hesperadin for *Pf*Ark1 over both *Pf*Ark2 and mammalian AurA and B.

### Inhibition of *Pf*Ark1 affects the progression of schizogony

We subsequently evaluated the effects of hesperadin and TAE684 against the proliferative stages of the parasites and their specificity for *Pf*Ark1. The rate at which hesperadin and TAE684 kill the parasite was evaluated by determining shifts in IC_50_ over time^45^. Hesperadin and TAE684 kill kinetics indicate that their effect is only evident with a significant IC_50_ shift after 48 h, indicative of activity within one life cycle (Figure 5A). This profile resembles that seen for the *Pf*PKG inhibitor ML10, which prevents parasite egress and invasion, but not for fast-acting compounds such as CQ (Figure 5A). To further evaluate which developmental stage during asexual proliferation is affected by hesperadin and TAE684 treatment, we treated tightly synchronized parasites at 12 h intervals, correlating to ring, early and late trophozoites and schizonts (Figure 5B). Hesperadin treatment had minimal impact on rings or trophozoites and did not limit the maturation from rings to trophozoites, whereas TAE684 treatment consistently affected trophozoite food vacuole formation (Figure 5B and 3C). The most pronounced effect for both compounds was associated with the completion of schizogony at ∼36–44 hpi, with no progression into the next cycle. Further evaluation of the morphologically aberrant schizonts revealed a significant reduction in the number of daughter merozoites formed for both hesperadin and TAE684 treatment (*p*<0.000001, n=30, unpaired Student’s t-test), an effect that was pronounced for the comparative treatment with TAE684 (Figure 5C). Hesperadin treatment did not prevent already formed schizonts from invading and forming rings in the next cycle, a process that was somewhat delayed by TAE684. Although this phenotype is reminiscent of the effect of *Pf*PKG inhibition, which also prevents parasite egress and subsequent invasion^46^, the lack of *Pf*PKG inhibition for hesperadin and TAE684 and their indifferent effect when combined with ML10 (ΣFIC_50_ of 1.1 and 1.3, respectively, Supplementary Figure S6) support a *Pf*PKG-independent mechanism for these compounds. However, the data delineate a timeframe of action of hesperadin associated with the completion of late schizogony processes.

**Figure 5:**
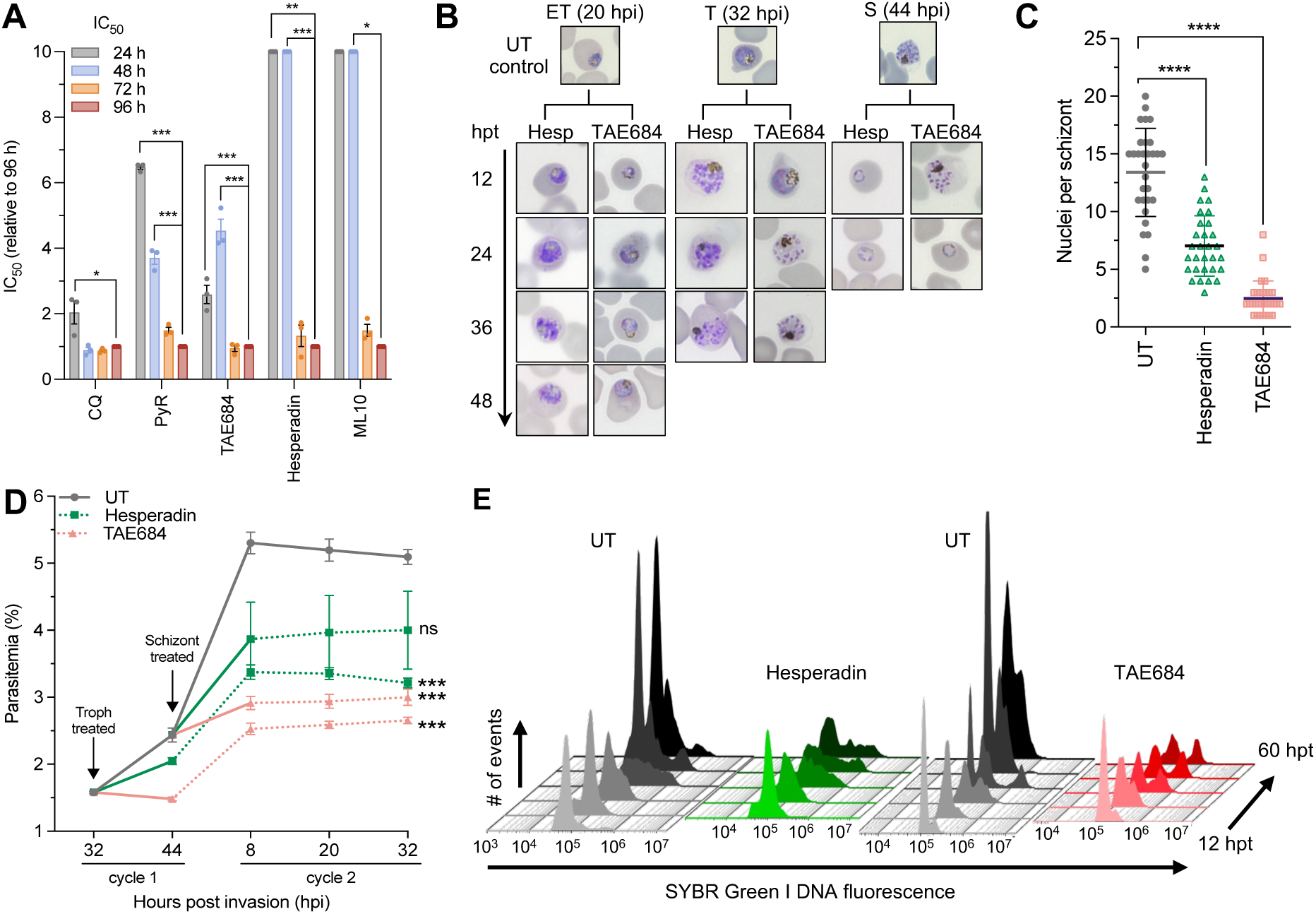
Effect of *Pf*Ark1 inhibition on intra-erythrocytic development. **A)** IC_50_ speed of kill assay using unsynchronised *Pf*NF54. Chloroquine (CQ), pyrimethamine (PyR). **B)** Morphological evaluation (Giemsa-stained thin smears) of *P. falciparum* asexual development following treatment with TAE684 (3xIC_50_, ∼900 nM) and hesperadin (IC_99_, ∼3 µM) (hpt – hours post-treatment; hpi – hours post-invasion). **C)** Nuclei count per schizonts after treatment (n=30). Error bars represent the 95% confidence interval (CI) of the mean. **D)** Synchronised *Pf*NF54 mature trophozoite and schizont populations treated with TAE684 and hesperadin for a 12 h period (solid line), washed off and parasitaemia measured at 12 h intervals using flow cytometry. **E)** Flow cytometric analysis of nuclear division following hesperadin and TAE684 treatment, sampled 12 hpt, and each subsequent 12 h until 60 hpt. Nuclei content was detected by consecutive staining with SYBR Green I (DNA fluorescence), detected in the FITC channel. Histograms overlaid for a representative sample of biological triplicates. All data are from three independent biological replicates, each performed in technical duplicates (n=3, mean ± S.E.), significance was calculated using a two-tailed Student’s *t*-test, \**p* <0.05, \*\*\**p* <0.001, \*\*\*\**p* <0.0001.

Flow cytometric quantification of this effect was performed after a 12 h drug treatment pulse on trophozoite and early schizont populations before drug washout (Figure 5D). Both Hesperadin and TAE684-treated trophozoites (∼30 hpi) were able to recover from a 12 h pulse and progress to schizonts, however, these parasites were unable to sufficiently establish reinvasion, resulting in a significant (*p*=0.00015 and *p*=0.00004, respectively, n=3, unpaired Student’s t-test) decrease in parasitemia in the subsequent population. A similar significant effect was immediately evident with TAE684-treated schizonts (*p*=0.0002, n=3, unpaired Student’s t-test) (∼40–42 hpi), however, no significant effect was observed for hesperadin-treated schizonts (*p*=0.139, n=3, unpaired Student’s t-test), reminiscent of the phenotype observed in schizont-treated samples (Figure 5B and 5D).

Quantification of the nuclear content of treated parasites indicated entry into schizogony comparative to untreated populations, but with the treatment primarily affecting parasites during schizogony and the fraction of individual cells halted in the schizont stage containing ≥4n (DNA content) was higher in drug-treated compared to untreated parasites (Figure 5E). Taken together, hesperadin as a *Pf*Ark1 inhibitor primarily affects parasite progression through schizogony (mid-to-late schizont development), where cells are unable to complete nuclear division and segregation successfully. However, it is ineffective against mature schizont stages, where these processes have already been completed.

### *Pf*Ark1 is critical to the completion of mitotic processes

Fluorescence microscopy morphological evaluation of ABS parasites, where *Pf*Ark1 activity was inhibited by hesperadin or TAE684, revealed nuclear morphological abnormalities in schizonts. The nuclei of those treated with hesperadin appeared distinctly multi-lobed, whereas TAE684-treated schizonts had more elongated nuclei (Figure 6A), although packaging into daughter cell structures did occur, as evident by nuclear membrane detection (Supplementary Figure S7A). The abnormal shape and decreased number observed during nuclear division were associated with abnormalities in MT structures in hesperadin– and TAE684-treated schizonts. Compared to the well-defined SPMTs seen in untreated schizonts, hesperadin treatment resulted in disorganized and interconnected MT structures. TAE684 decreased the appearance of MT structures entirely, with an overall decrease in nuclear content (Figure 6A).

**Figure 6:**
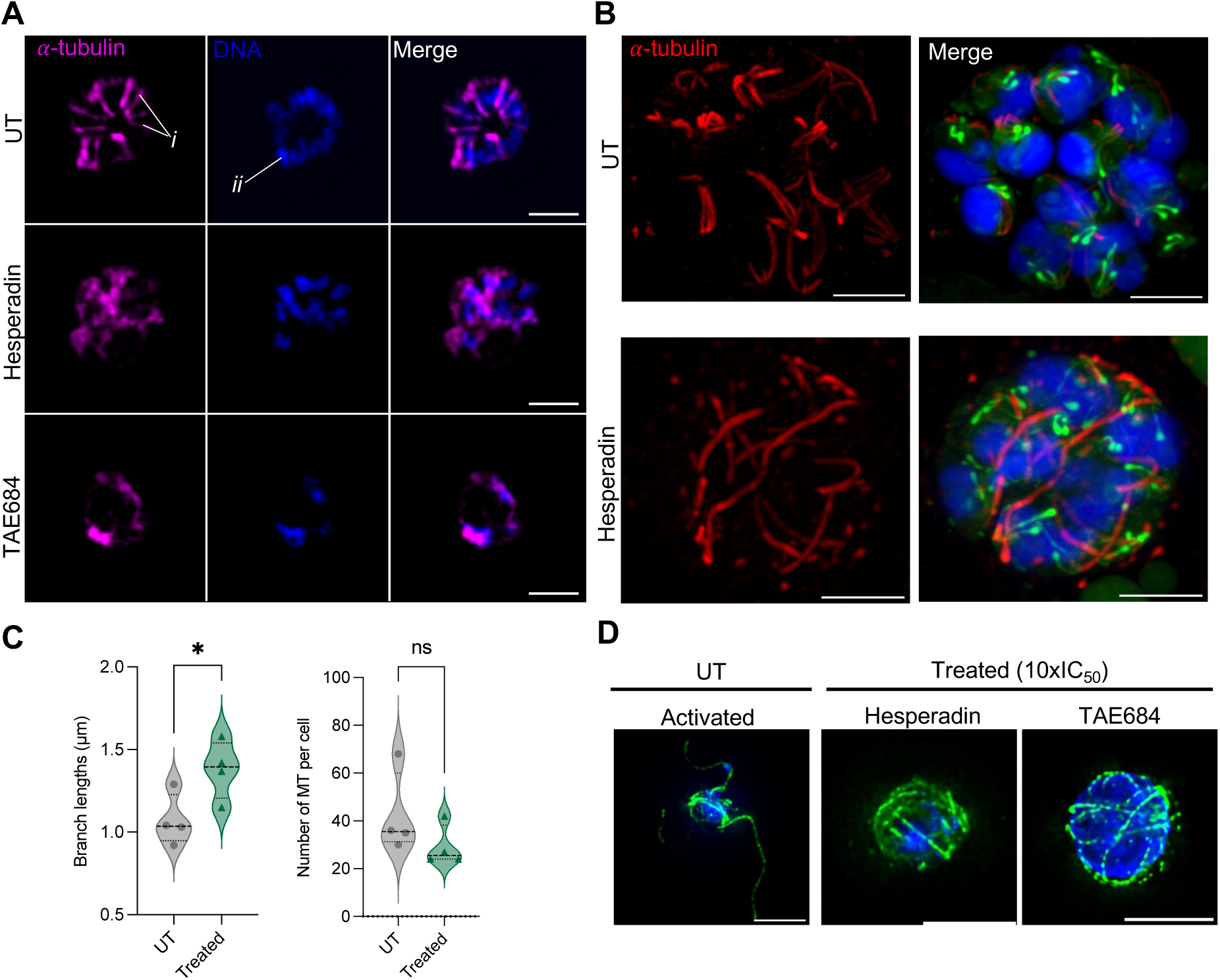
*Pf*Ark1 inhibition effect on microtubule and nuclear material morphology. Ring stage, synchronised, asexual intra-erythrocytic *Pf*NF54 parasites treated with hesperadin and TAE684 were harvested at ∼46 hpi. **A)** Representative images (maximum intensity projections) of the morphological abnormalities observed in nuclei (Hoechst, blue) & microtubules (anti-⍺-tubulin, pink), showing subpellicular microtubules (SPMTs, *i*), nuclei (*ii*). The images represent at least ten parasites per sample. Scale bars correspond to 2 µm. **B)** Representative images (maximum intensity projections) of expansion microscopy images of UT mature schizonts (top panel) compared to hesperadin-treated (bottom panel). Red = tubulin, blue = DNA, and green = NHS-ester. Scale bars correspond to 5 µm. **C)** Average microtubule (MT) branch lengths and number of MT branches per cell of UT compared to hesperadin-treated, n=4 schizonts, mean ± SD with an unpaired Welch’s two-tailed t-test. **D)** Representative images of TAE684 and hesperadin-treated *P. falciparum* male gametes labelled with anti-⍺-tubulin (green) and co-stained with DAPI for nuclei (blue). Scale bars correspond to 5 µm.

Expansion microscopy (ExM) was subsequently used to provide more nuanced evaluation of hesperadin-treated cells, which revealed the extent of abrogation of nuclear segmentation and packaging, as well as extensive MT defects (Figure 6B). Untreated parasites contained the expected well-packaged and segregated daughter nuclei, with SPMTs clearly extending from the MTOCs associated with the CPs in proximity to well-formed rhoptries (Figure 6B). No intranuclear MTs were evident as segregation was completed. By contrast, hesperadin treatment caused parasites to halt in schizogony at a point where the parasites contained multilobed nuclei, with the nuclear material not separated in most instances, and nuclei showing an increased nuclear volume. The MT organization was clearly affected by hesperadin treatment, with significantly extended MT structures (average lengths of 1.4 ± 0.7 µm, *vs*. untreated parasites at 1.1 ± 0.4 µm, *p=*0.04, n=4, Figure 6C) spanning across and connecting different nuclear centers/CPs. Fewer of these MT structures were present per cell (29 *vs*. 42 in hesperadin *vs*. UT cells), and fewer could be associated with MTOC and the limited rhoptry pairs formed. This suggests that these microtubule structures are halted as interpolar MTs that are unable to retract during the interpolar to hemi-spindle transition, as is typically required. ^47,48^ This suggests that *Pf*Ark1 function is necessary for meta– and anaphase-like transition processes during mitosis in the parasite. However, the potential for these aberrant MTs to be malformed, extended SPMTs cannot currently be excluded. Indeed, some of these aberrant MT structures closely localize to rhoptries connected to associated apical polar rings, as one would expect from SPMTs (Figure 6B).

We subsequently evaluated the broader involvement of *Pf*Ark1 in mitotic processes also present during male gametogenesis. Under normal conditions, when mature male gametocytes are successfully activated, exflagellation results in the formation of eight MT-labelled flagella, each associated with segregated nuclei (Figure 6D) compared to non-activated mature gametocytes (Supplementary Figure S7B). Treatment with TAE684 and hesperadin was associated with some DNA replication proceeding normally, but the nuclear material remained highly compacted and lacked proper segregation. Strikingly, the MT structures in the drug-treated cells are abnormal, thin and wrapped around the nucleus, pointing to disorganized MT forming flagellae (Figure 6D). This suggests that DNA replication has mostly occurred, but the absence of *Pf*Ark1-mediated signaling disrupted microtubule organization on the basal body in these stages. The extended, thin phenotype closely resembled that observed during late schizogony under *Pf*Ark1 inhibition (Figure 6B). These findings indicate that *Pf*Ark1 activity is required post-DNA replication but before nuclear segregation and egress of male gametes, suggesting a mitotic block during gametogenesis^36^. These findings suggest that the disruption of *Pf*Ark1 function during male gametogenesis does not affect DNA replication but rather may disrupt proper kinetochore attachment and spindle formation.

## Discussion

Several kinases are validated as drug targets in *P. falciparum*, and this still presents an enticing family of proteins to target in this organism for the development of antimalarial candidates. In this work, we expand the current knowledge base regarding kinases as drug targets in malaria parasites by chemically validating the mitotic kinase *Pf*Ark1 as an antimalarial target. To our knowledge, this is the first direct chemical evidence of an inhibitor specifically targeting any of the *Pf*Ark members. Notably, we demonstrate that inhibition of *Pf*Ark1 activity can be achieved at single-digit nanomolar concentrations, with a large selectivity window for the parasite, and a favorable profile for potential drug candidates.

We demonstrate that human Aurora kinase inhibitors can specifically inhibit all proliferative stages of *P. falciparum* parasite development, while also displaying selectivity to the parasite. Our data show that *Pf*Ark1 is a key player amongst the Ark family members and can be inhibited to prevent the proliferation of ABS parasites, hepatic schizogony and male gamete exflagellation. In addition to having potential as ABS active in TCP-1 type strategies^49^, this raises the possibility of leveraging *Pf*Ark1 inhibition in transmission-blocking strategies (TCP-5 potential)^50^. Even though both *Pf*Ark1 and *Pf*Ark2 are expressed during gametocytogenesis^35^, their functional impairment is only evident during male gametocytogenesis, similar to other gametocyte-sterilizing compounds^51^.

The three most active inhibitors, hesperadin, TAE684, and AT83, all displayed no cross-resistance to known antimalarial-resistant parasite lines, suggesting that each has a unique mode of action against *P. falciparum* asexual parasites. We then verified, through various biochemical methods, that both hesperadin and TAE684 were *Pf*Ark1 inhibitors, with hesperadin specifically and selectively inhibiting *Pf*Ark1. TAE684 displayed additional polypharmacological characteristics, including hemozoin formation inhibition, similar to other kinase inhibitors^52^. To our knowledge, our data provide the first indication of anti*plasmodium* activity of AT83, a compound that has advanced in multiple early-phase clinical trials with favorable toxicity profiles^53^. These data expand and diversify the compendium of targetable mitotic kinases in *Plasmodium* beyond the current indication of *Pf*Nek3 inhibition with BI-2536, a known potent human polo-like kinase 1 inhibitor^54^. This provides a clear starting point for drug repurposing and repositioning strategies against malaria. Hesperadin is also active against *T. brucei* and *Leishmania major,* with initial hit expansion indicating that analogues mirror hesperadin’s activity and phenotype^24^. This provides support for SAR expansion studies and further development of hesperadin as an anti*plasmodium* chemotype.

In this study, we demonstrate that *Pf*Ark1 is the most vulnerable target from the Ark protein family in *P. falciparum*, correlating with its essential requirement in ABS parasite^10^. Moreover, *Pf*Ark1 inhibition abrogates mitotic-associated proliferation, suggesting that *Pf*Ark1 is the primary aurora kinase member governing mitotic-related processes^14^ in ABS parasites, male gametes^55^ and most likely also during hepatocyte schizogony. With hesperadin as a tool compound that exclusively inhibits *Pf*Ark1, we could deduce the functional and mechanistic importance of this protein during mitosis. *Pf*Ark1 inhibition results in abnormal nuclear morphology, along with spindle structure defects that we previously described^27^.

Our high-resolution imaging provides the first evidence that disruption of *Pf*Ark1 function causes complete disorganization of MT structures required to coordinate and complete mitosis. The multilobed nuclei observed are reminiscent of the inhibition of AurB (equatorial aurora kinase) in other organisms and apicomplexan parasites, which results in misaligned chromosomes, lagging chromatids, cytokinesis failure, and polyploidy^26,30,56^. Our observations support *Pf*Ark1 to be functionally similar to AurB in its association with kinetochores until metaphase, and the extended MT structures after loss of *Pf*Ark1 function imply a lack of translocation to the central spindle to coordinate cytokinesis. Although the disruption of AurB function can lead to disruption of microtubule dynamics and structure, these effects are also observed in the inhibition of AurA, which leads to unaligned chromosomes due to impaired centrosome separation and the formation of monopolar spindles, which could explain the spindle structures we observe in mature treated schizonts^19,57^. Thus, based on these phenotypic observations, we suggest that *Pf*Ark1 fulfils the mitotic responsibilities of both AurA and AurB to control correct mitotic and interpolar spindle formation in ABS parasites and male gametocytes. Co-localization with known markers of the centrosome (e.g., Centrins^58^), kinetochores (e.g., NDC80^59^) and chromosome passenger proteins (e.g., INCENP), will be needed to confirm if *Pf*Ark1 takes on a role of functional redundancy in *P. falciparum* for both AurA and B.

This study therefore describes hesperadin as the first potent and selective inhibitor of *Pf*Ark1 in *Plasmodium falciparum*, exhibiting activity across multiple life cycle stages of the parasite that can be considered for future development as an antimalarial. Additionally, we pose that *Pf*Ark1 is the most important aurora kinase regulating mitotic processes in the parasite.

## Methods

Parasitology work and volunteer human blood donation (from consenting, healthy adult volunteers) at the University of Pretoria is covered under ethical approval from the Health Sciences Ethics Committee (506/2018) and Natural and Agricultural Sciences Ethics Committee (180000094). Full methods are provided in the supplementary material.

### Parasite *in vitro* cultivation

*P. falciparum* drug sensitive NF54 (*Pf*NF54) and drug-resistant Dd2 (*Pf*Dd2, chloroquine, pyrimethamine and mefloquine resistant) strains were cultured *in vitro* in human O^+^/A^+^ erythrocytes (5% hematocrit) in complete medium at 2–5% parasitemia, 37 °C, shaking at 60 rpm under hypoxic conditions (90% N_2_, 5% O_2_ and 5% CO_2_). Immature and mature gametocytes were produced from *P. falciparum* NF54cg6-ULG8-CGB99^60^ as described^61,62^ by simultaneously applying nutrient starvation and decreasing hematocrit to induce >97% ring-stage ABS parasites (0.5% parasitemia, 6% hematocrit). Gametocytogenesis was followed with daily media changes (including glucose) and ABS parasites removed with 50 mM N-acetylglucosamine (day 1–4 for stage II/III; days 3–7 for stage IV/V).

### *In vitro* asexual and gametocyte activity evaluation

All inhibitors were dissolved in dimethyl sulfoxide (DMSO). Anti*plasmodium* activity of selected anti-cancer inhibitors was determined on *P. falciparum* NF54 and Dd2 strains, measuring SYBR Green I fluorescence (485 nm excitation, 538 nm emission) as an indicator of proliferation^63^. Ring-stage ABS parasites (1% parasitemia, 1% hematocrit) were treated with compounds for 96 h at 37 °C with chloroquine disulphate (0.5 μM) as positive drug control. The activity of the inhibitors was tested against immature (>95% stage II/III) and late-stage (>90% stage IV/V) gametocytes using a luciferase reporter assay on NF54cg6-ULG8-CGB99. Cultures (2% gametocytemia, 1.5% hematocrit) was exposed to drug pressure for 48 h with methylene blue (5 μM) as positive control for inhibition. Luciferase activity was measured in 20 μL parasite lysates by adding 50 μL luciferin substrate (Promega Luciferase Assay System) at room temperature, detecting bioluminescence at an integration constant of 10 s. Assay performance was monitored with Z-factors > 0.8.

### *P. berghei* (PbLuc) liver stage screening

Liver stage (PbLuc) screening has been previously described^64^. *Plasmodium berghei* (*Pb*) sporozoites were obtained by dissecting the salivary glands of infected *Anopheles stephensi* mosquitoes, provided by The SporoCore, at the University of Georgia, GA, USA (<u>SporoCore.uga.edu</u>). Engineered parasites, termed Pb-Luc (GFP-Luc_ama1-eef1_ reporter line^65^) were used to infect HepG2-A16-CD81EGFP cells, stably transformed to express a GFP-CD81 fusion protein^66^. 3 × 10^3^ cells were seeded in 1536-well plates, with atovaquone and puromycin as positive inhibition controls, and 0.1% DMSO for negative inhibition. The following day, Pb-Luc sporozoites were extracted and purified from the salivary glands of infected mosquitoes, 5 μL of which were added to the cell plates at 1 × 10^3^ sporozoites per well and incubated at 37 °C and 5% CO_2_ for 48 h. Luciferase reporter gene expression was quantified by luminescence using the Bright-Glo Luciferase Assay System on a PHERAstar FSX.

### *P. falciparum* Dual Gamete Formation Assay (*Pf* DGFA)

The *Pf* DGFA^67^ was performed by incubating *Pf*NF54 gametocytes with test molecules in 384-well plates for 48 h. Gametogenesis was triggered with the addition of xanthurenic acid and a temperature decrease. Male and female gametes were identified through fluorescent microscopy and quantified from recorded data using a custom imaging algorithm and then compared to negative (DMSO) and positive (1 µM cabamiquine) controls. Percentage inhibition was calculated, and dose-response curves were constructed in GraphPad Prism v10.5 to calculate IC_50_ values.

### Cytotoxicity counter-screening

Maintenance and cytotoxicity screening of hepatocellular carcinoma (HepG2) and Chinese ovarian cells (CHO) were conducted as before^68^. Cells were seeded at 1×10^4^ cells/well in a 96-well plate and incubated for 24 h, then treated in duplicate with a 10-fold serial dilution of the inhibitor, starting at 100 μg/mL, in new media and incubated for 48 h. HepG2 and CHO toxicity measurements were performed using the MTT tetrazolium reduction assay and absorbance at 540 nm (Multiskan GO). Emetine was included as a positive drug control.

### Rate– and stage-specific evaluations and inhibitor reversibility

Tightly synchronized *Pf*NF54 cultures were treated with 3xIC_50_ of TAE684, AT83, and ZM-39 and IC_99_ for hesperadin at ring, trophozoite, and schizont stages and sampled every 12 h over a 48 h period. Parasites were monitored morphologically using Giemsa-stained thin smears. Images were captured using a Nikon Eclipse 50i light microscope adapted with a Nikon camera (Nikon DS-Fi1) and NIS-Elements software. The IC_50_ speed assay was conducted as described^45^. To determine whether the effect is reversible for each inhibitor, synchronized late trophozoite and schizont stage parasite cultures were treated with TAE684 at 3xIC_50_ and hesperadin at IC_99_ for 12 h, after which the inhibitor was washed off and parasite progression monitored every 12 h over a 48 h period using flow cytometry^15^.

### Inhibitor susceptibility assays using *P. falciparum* kinase cKD lines

Compound susceptibility assays using *P. falciparum* Ark1 and Ark2 cKD lines were performed^69^ on synchronous ring-stage Ark1 (PF3D7_0605300) and Ark2 (PF3D7_0309200) cKD parasites (see supplementary methods), controlled with a parasite line expressing an aptamer-regulatable fluorescent protein, maintained in the presence of high anhydrotetracycline (aTc, 500 nM) or no aTc in 384-well microplates. Serially diluted compounds, DMSO and dihydroartemisinin treatment (500 nM) were included and incubated for 72 h. Luminescence was measured using the Renilla-Glo Luciferase Assay System on a GloMax Discover Multimode Microplate Reader, and IC_50_ values were obtained from corrected dose-response curves using Graph-Pad Prism v10.5.

### Antimalarial resistome barcode sequencing (AReBar) cross-resistance assay

Cross-resistance of Hesperadin, TAE684 and AT9283 was evaluated against a pool of barcoded drug-resistant parasites^39^. The pool consisted of 52 lines (Supplementary File S2) in both the Dd2 and 3D7 backgrounds, including wild-type, that were barcoded at the *pfpare* locus (PF3D7_0709700)^70^, with unique 11-bp barcode sequences. The pool (triplicate cultures of 1 mL) was exposed to 3 x IC_50_ of each test compound, including the positive control compound halofuginone that targets prolyl tRNA synthetase. Growth of the pool was measured every 2–3 days by flow cytometry (Beckman CytoFlex 5), staining with 1xSYBR Green I and 200 nM MitoTracker Deep Red, and parasitemia maintained within a range <5% parasitemia. At day 14, samples were harvested, lysed with 0.05% saponin and parasite pellets collected. Barcodes were amplified by PCR and quantified by sequencing using an Oxford Nanopore minION. The change in barcode proportion (log_2_ fold change, LFC) was measured relative to the no-drug control, with LFC>2.5 indicating cross-resistance.

### *Pf*PKG, *Pv*PI4Kβ and *Pf*CLK3 Kinase Assays

Full-length *Pf*PKG (PF3D7_1436600)^22^ and *Pv*PI4Kβ (PVX_098050)^71^ recombinant proteins were expressed in *E. coli* and baculovirus-insect cell expression systems. The kinase domain of *Pf*CLK3 (PF3D7_1114700, Gln317 to Ser694) was expressed in *E. coli* Rosetta™(DE3)pLysS cells (kind gift from Rafael M. Couñago, CQMED, University of Campinas). Briefly, the N-terminal His-tagged kinases were purified using immobilized metal affinity chromatography, followed by anion exchange chromatography in the case of *Pf*PKG, and size exclusion chromatography. Dose-response kinase inhibition assays were carried out using the ADP-Glo kinase assay kit^72^, with 1 nM *Pf*PKG, 10 μM ATP, 20 μM GRTGRRNSI-NH_2_, 1% (v/v) DMSO and inhibitor in PKG assay buffer (25 mM HEPES pH 7.4, 0.1 mg/mL BSA, 0.01% (v/v) Triton-X 100, 20 mM MgCl_2_, 2 mM DTT, 10 μM cGMP); or 6 nM *Pv*PI4Kβ, 10 μM ATP, 0.1 mg/ml L-alpha-phosphatidylinositol, 1% (v/v) DMSO and inhibitor in PI4Kβ assay buffer (25 mM HEPES pH 7.4, 100 mM NaCl, 3 mM MgCl2, 1 mM DTT, 0.025 mg/ml BSA, 0.2% (v/v) Triton-X-100); or 15 nM *Pf*CLK3, 10 μM ATP, 20 μM myelin basic protein, 1% (v/v) DMSO and inhibitor in assay buffer (50 mM HEPES pH 7.4, 0.1 mg/mL BSA, 0.01% (v/v) Triton-X 100, 20 mM MgCl2, 2 mM DTT). The data were normalized based on the 100% activity controls (1% DMSO) and the 100% inhibition controls [ML10 for *Pf*PKG, MLN0128 (sapanisertib^72^) for *Pv*PI4Kβ and TCMDC-135051^73^ for *Pf*CLK3, at 10 µM]. Mean IC_50_ values were calculated from two independent experiments, each with technical duplicates.

### Molecular docking

A homology model for *P. falciparum* Ark1 (*Pf*Ark1) was generated using the crystal structure of human Aurora kinase A (*Hs*AurA) co-crystallized with ATP (5DNR)^74^, with a 33% sequence identity. Ligands (LigPrep tool, pH 7.4 ± 0.5, OPLS4 forcefield) and the protein structures were prepared, analyzed and visualized in the Schrödinger 2021-4 Maestro interface. After structural optimization, the receptor grid was generated with 10 Å inner & 20 Å outer grid boxes over hinge region residues Phe107-The119. Ligands were docked to the receptor using the Glide default settings in extra precision, flexible ligand mode, and poses were scored using the Schrödinger GlideScore function^75^. Analysis of the docking poses was conducted using Molecular Mechanics and Generalized Born Surface Area (MM-GBSA) with the default solvation model (VSGB and OPLS4) settings, with flexible residues within 20 Å of the active site.

### Isobologram analysis

Fixed-ratio isobologram analysis^76^ was performed on parasites were treated with fixed ratios (5:0, 4:1, 3:2, 2:3, 1:4, and 0:5) of TAE684 and hesperadin in combination with chloroquine (CQ), ML10, and each other. Similarly, this was also done for AT83 and ZM-39 in combination with CQ and each other using the SYBR Green I proliferative assay as readouts to determine the fractional inhibitory concentrations (FIC) for the respective combinations. Isobolograms were generated by plotting paired FIC values linearly, utilizing the average of three biological replicates, each performed in technical duplicates. The paired FIC values for each drug combination were analyzed, and the mean FIC values (ΣFIC) were calculated to delineate synergism (<0.8), additivity/indifference (0.8–1.4), or antagonism (>1.4).

### NP-40 mediated cell-free β-hematin inhibition assay

The β-hematin inhibition assay described in^77^ was used to test the inhibitors for their ability to inhibit β-hematin formation. Briefly, serial dilutions of each inhibitor in an “NP-40 substitute” detergent (305.5 μM) were tested at a starting in-well concentration of 1000 μM or 500 μM. 25 mM hematin was added, and plates were incubated for 5 h at 37 °C, after which 32 μL of a 50% pyridine solution and 60 μL of acetone were added to all wells. Heme-pyridine complex was detected by absorbance at 405 nm. Heme fractionation analysis entailed exposing ring-stage NF54 parasites to compounds at multiples of their ABS IC_50_. After a 32 h incubation, parasites were isolated with saponin and trophozoites were subjected to a series of lysis, solubilization, and differential centrifugation steps to obtain fractions that contained Hb, free heme or Hz^43^. The UV-visible spectra of heme as Fe(III)heme-pyridine were quantified, normalized to the number of analyzed cells as determined via flow cytometry.

### Fluorescence and U-ExM microscopy

*In vitro*, synchronized ring-stage parasite cultures were treated at TAE684 (3xIC_50_) and hesperadin (IC_99_) and sampled at ∼36 h post-treatment (∼46 hpi). Nuclear content and microtubule morphology were assessed using direct immunofluorescence imaging on fixed parasites (4% PFA, 15 min) on poly-D-lysine-coated coverslips. These were washed three times with PBS, permeabilized with fresh 0.1% Triton X-100, washed another three times with PBS, and blocked with 3% BSA-PBS for 1 h at room temperature (RT). Coverslips were then exposed to 1:500 dilution of primary anti-tubulin, mouse monoclonal (Merck, T5192), overnight at 4 °C. Primary antibody was subsequently washed off three times with PBS and incubated in anti-mouse conjugated CF488A secondary antibody produced in chicken (Merck, SAB4600238) for 1 h at RT. Cells were then mounted on a glass slide using ProLong™ glass antifade mount medium with NucBlue™ (P36981, ThermoFisher). Images were acquired using a Zeiss LSM 780 Inverted Confocal Laser Scanning Microscope (LSM) (Zeiss, Germany) for super-resolution imaging in the appropriate channels with a ×100 oil-immersion objective and 1.4 numerical aperture. Images were processed using Zeiss ZEN Lite Blue Edition software (Zeiss, Germany) and Fiji software. For ultra-expansion microscopy (U-ExM), the procedure was performed as previously described in^48^. Images were acquired using an LSM 980 with Airyscan 2 (Zeiss, Germany) for super-resolution imaging in the appropriate channels with a ×100 oil-immersion objective and 1.4 numerical aperture. Images were processed using Zeiss ZEN Lite Blue Edition software (Zeiss, Germany) and Fiji software. Experimental and imaging analysis pipeline details are included in the supplementary methods.

### Statistical analysis

A two-tailed t-test (95% CI) was used for the statistical assessment of measurement differences that might present significance relative to the control. The significant differences are displayed as asterisks on graphs: * *p* < 0.05; ** *p* < 0.01; *** *p* < 0.001, \*\*\*\**p* < 0.0001. The data presented was from one experiment performed in quadruplicate and subsequently analyzed using Microsoft Excel and GraphPad Prism v10.5 software.

## *Corresponding Author

Lyn-Marie Birkholtz

## Author Contributions

HL performed the work with MM, T Rabie and JT (modelling and docking). HL and KM performed the hematin experiments. KM performed the hemozoin experiments under KJW, with NS performing recombinant protein work under LBC with KC. SGD, LCG, NB, MTF, MLdS, JF contributed experimental data with interpretations and validations under JCN, MSL, MD and EAW. LMB conceptualized the study and wrote the paper with HL3. All co-authors contributed to and approved the final version of the manuscript.

## Acknowledgements

We thank Ben Liffner and Sabrina Absalon (Indiana University) for advice on U-ExM and Mike Reiche at the African Microscopy Initiative for microscopy support. We thank Reena Zutshi from Luceome Biotechnologies for the KinaseSeeker™ assays. This project was in part supported by the Medicines for Malaria Venture (LMB: RD-19-001), the South African Medical Research Council (KC) and the Department of Science and Innovation South African Research Chairs Initiative Grants managed by the National Research Foundation (LMB UID: 84627). The University of Pretoria Institute for Sustainable Malaria Control acknowledges the South African Medical Research Council as Collaborating Centre for Malaria Research. K.C. is the Neville Isdell Chair in African-centric Drug Discovery and Development and thanks Neville Isdell for generously funding the Chair. MTF is supported by a grant to MJD awarded by the Medicines for Malaria Venture (RD-21-1003). MJD is supported by a UKRI MRC Career Development Award (MR/V010034/1).

## Abbreviations

ABS: Asexual blood stage
Aur: Aurora kinase
AReBar: Antimalarial resistome barcode sequencing
βH: β-hematin
CQ: chloroquine
cKD: conditional knockdown
CP: centriolar plaque
MT: microtubule
MTOC: microtubule-organizing center
hpi: h post invasion
hpt: h post treatment
U-ExM: ultra-expansion microscopy.

