## Supplementary methods and figures for "Targeting Aurora kinases as essential cell cycle regulators to deliver multi-stage antimalarials against *Plasmodium falciparum*"

**Parasite *in vitro* cultivation**

*P. falciparum* drug sensitive NF54 (*Pf*NF54) and drug-resistant Dd2 (*Pf*Dd2, chloroquine, pyrimethamine and mefloquine resistant) strains were cultured *in vitro* in human erythrocytes (5% hematocrit) in complete medium [RPMI-1640 with 23.81 mM sodium bicarbonate, pH of 7.4; 0.5% (w/v) Albumax II, 80 mg/mL gentamycin, 25 mM HEPES, 20 mM glucose, and 0.2 mM hypoxanthine] at 2–5% parasitaemia, 37 °C, shaking at 60 rpm under hypoxic conditions (90% $N_2$, 5% $O_2$ and 5% $CO_2$). The medium was replaced daily, and parasite morphology and proliferation were monitored by Giemsa-stained slides visualised by light microscopy using an x100 oil immersion lens at x1,000 magnification. Immature and mature gametocytes were produced from *P. falciparum* NF54cg6-ULG8-CGB99[1] as described[2,3] by simultaneously applying nutrient starvation and decreasing hematocrit to induce >97% ring-stage ABS parasites (0.5% parasitemia, 6% hematocrit). Gametocytogenesis was followed with daily media changes (including glucose) and ABS parasites removed with 50 mM N-acetylglucosamine (day 1-4 for stage II/III; days 3-7 for stage IV/V).

***In vitro* intra-erythrocytic asexual screening**

All inhibitors were dissolved in dimethyl sulfoxide (DMSO). Antiplasmodial activity of selected anti-cancer inhibitors was measured on *P. falciparum* NF54 and Dd2 strains using a SYBR Green I proliferation-based assay as described in[4]. Briefly, inhibitors were serially diluted in a 96-well plate, and the parasite suspension (1% parasitaemia, 2% hematocrit) was subsequently added to controls consisting of no-treatment wells containing drug-free media. In contrast, no-grow control wells contained 50 µM chloroquine (CQ). The plates were then incubated at 37 °C for 96 h in a low-oxygen atmosphere, after which parasite proliferation was measured using 10x Syber Green I (Invitrogen) diluted (1:5000) in lysis buffer (20 mM Tris/HCl, 5 mM EDTA, 0.16% (w/v) saponin, 1.6% (v/v) Triton X). The plates were incubated at room temperature (RT) for 45–60 min. Parasite proliferation was assessed by measuring the fluorescence signal (485 nm excitation, 530 nm emission) using Fluoroskan Ascentt FL (Thermo Labsystems), and $IC_{50}$ values were determined using GraphPad Prism v10.5, where the medium containing background was subtracted.

***P. berghei* liver stage screening**

*Plasmodium berghei* (Pb) sporozoites were obtained by dissecting the salivary glands of infected *Anopheles stephensi* mosquitoes. These were provided by The SporoCore, at the University of Georgia, GA, USA (SporoCore.uga.edu). The parasites utilised a GFP-$Luc_{ama1\text{-}eef1}$ reporter line[5]. These engineered parasites, termed Pb-Luc, were used to infect HepG2-A16-CD81EGFP cells. HepG2-A16-CD81EGFP cells stably transformed to express a GFP-CD81 fusion protein[4] were cultured at 37 °C in 5% $CO_2$ in DMEM (Invitrogen, Carlsbad, USA) supplemented with 10% FBS (Corning, NY, USA), 1X Pen/Strep/Glu (Thermo Fisher Scientific, USA).

Compounds were prepared as 10 mM solutions in DMSO, and 10 nL of each (resulting in a final DMSO concentration of 0.1% per well) was transferred into assay plates using an ECHO 650 (Beckman). Concentrations ranged from 10 µM to 0.5 nM. Atovaquone (0.1 µM) served as a positive control, while 0.1% DMSO was used as a negative control.

Liver stage (PbLuc) screening has been previously described[6]. Human hepatic cells ($3 \times 10^3$; HepG2-A16-CD81-EGFP) suspended in 5 µL of DMEM medium ($2\times10^5$ cells/mL, supplemented with 5% FBS, 5× Pen/Strep/Glu) were seeded in 1536-well plates (Greiner BioOne) using the MultiFlo FX (Agilent) 20 h before infection. PbLuc sporozoites, isolated from the salivary glands of *A. stephensi* mosquitoes, were filtered twice through a 20 µm nylon pore cell strainer. The sporozoites were resuspended in screening media, counted with a hemocytometer, and diluted to a final concentration of 200 sporozoites per µL. Each well received 1,000 sporozoites in 5 µL, dispensed via the GNF Dispenser II (GNF Systems). The plates were then centrifuged for 3 min in an Eppendorf 5810 R (330 RCF) at the lowest acceleration and brake settings. After incubation at 37 °C with 5% $CO_2$ for 48 h, the media was removed by centrifuging the inverted plates at 130 RCF for 1 min. Wells then received 2 µL of Bright-Glo™ Luciferase Assay System (Promega). Luminescence was measured immediately with the Pherastar FSX reader (BMG Labtech). Luminescence values were normalized using the positive and negative controls. $EC_{50}$ values were calculated with CDD Vault (Burlingame, CA). All experiments were performed with at least four technical replicates and repeated three times.

***In vitro* gametocyte screening**

All inhibitors were dissolved in dimethyl sulfoxide (DMSO). Antiplasmodial activity of selected anti-cancer inhibitors was measured on *P. falciparum* NF54 and Dd2 strains, measuring SYBR Green I fluorescence (485 nm excitation, 538 nm emission) as an indicator of proliferation[4]. Ring-stage ABS parasites (1% parasitemia, 1% hematocrit) was treated with compounds for 96 h at 37 °C with chloroquine disulphate (0.5 µM) as positive drug control. The activity of the inhibitors was tested against immature (>95% stage II/III) and late-stage (>90% stage IV/V) gametocytes using a luciferase reporter assay on NF54cg6-ULG8-CGB99[1,3,4]. Cultures (2% gametocytemia, 1.5% hematocrit) was exposed to drug pressure for 48 h with methylene blue (5 µM) as positive control for inhibition. Luciferase activity was measured in 20 µL parasite lysates by adding 50 µL luciferin substrate (Promega Luciferase Assay System) at room temperature and detection of resultant bioluminescence at an integration constant of 10 s. Assay performance was monitored with Z-factors at > 0.8.

**_P. falciparum_ Dual Gamete Formation Assay (_Pf_ DGFA)**

The *Pf* DGFA was performed as described[7]. Briefly, *P. falciparum* NF54 strain gametocytes were incubated with test molecules in 384 plates for 48 h. Then, gametogenesis was triggered with the addition of xanthurenic acid and temperature decrease from 37 °C to 4 °C for 4 min, then 28 °C for a further 5 min. 20 min after triggering gametogenesis, plates were imaged by timelapse brightfield microscopy to identify male gametogenesis by observing motile exflagellation centres. Then, plates were maintained at 28 °C for a further 24 h to allow the maximal expression of *Pf*s25 on the surface of female gametes. Female gametes were observed by live staining with anti-*Pf*s25 (clone 4B7, BEI Resources) coupled to Cy3 fluorophore and automated fluorescence microscopy. Male and female gametes were identified and quantified from recorded data using a custom imaging algorithm and then compared to negative (DMSO) and positive (1 µM cabamiquine) controls. Percentage inhibition was calculated, and dose response curves were constructed in GraphPad Prism v10.5 to calculate $IC_{50}$ values.

**Cytotoxicity counter-screening**

HepG2 and CHO cells were maintained at 37 °C, under a humidified atmosphere of 5% $CO_2$, in DMEM high glucose media (Gibco) supplemented with 10% heat-inactivated FBS and 1% (v/v) Penicillin-Streptomycin antibiotics. Cells were grown to 90% confluency, after which the cells were washed with PBS and trypsinised with trypsin-EDTA (0.25% Trypsin and 1 mM EDTA, Merck) and centrifuged for 2 min (10 000 x*g*). The supernatant was discarded, and the cell pellet was resuspended in 1 mL DMEM media for cell counting by Trypan Blue. To determine the toxicity, cells were resuspended and seeded at a concentration of $1x10^4$ cells/well in a 96-well plate and incubated for 24 h[8,9]. Cells were treated in duplicate with a 10-fold serial dilution starting at 100 ug/mL of inhibitor in new media. After 48 h incubation, the viable cells were measured using the MTT tetrazolium reduction assay, where sterile 25 µL of a 5 mg/mL MTT was added to each well and incubated for 4 h before the water-soluble formazan crystals were dissolved with 100 µL DMSO (Merck) per well. The absorbance, a measure of intracellular reduction of MTT, was measured at 540 nm (Multiskan GO). Emetine (50 µM) was included as a positive drug control, and the background was subtracted from all data before being normalised to the untreated[8,9].

**Rate- and stage-specific morphological evaluations and inhibitor effect reversibility**

To determine the stage specificity during asexual intra-erythrocytic development, *P. falciparum* NF54 parasites were synchronised using 5% D-Sorbitol to obtain a >90% ring-stage population (6 – 10 h postinvasion, hpi). In *vitro*, synchronised parasite cultures were treated at 3x$IC_{50}$ for TAE684, AT83, and ZM-39 and at $IC_{99}$ for hesperadin. The rate of activity and stage-specificity was monitored morphologically using Giemsa-stained thin smears. Asexual intra-erythrocytic stage parasites were treated at ring, early/late trophozoite, and schizont stages, sampling every 12 h over a 48-hour period. Images were captured using a Nikon Eclipse 50i light microscope adapted with a Nikon camera (Nikon DS-Fi1) and NIS-Elements software. To determine how fast the inhibitor affects a specific parasite stage and whether the effect is reversible for each inhibitor, synchronised late trophozoite and schizont stage parasite cultures were treated with TAE684 at 3x$IC_{50}$ and hesperadin at $IC_{99}$ for 12 h, after which the inhibitor was washed off with inhibitor-free medium. Parasite progression was monitored every 12 h over a 48-h period using flow cytometry, as described in[7]. The $IC_{50}$ speed assay was conducted as described in[8]. Briefly, unsynchronised *P. falciparum* NF54 asexual proliferation was determined using a SYBR Green I proliferation-based assay as described in the ref[2] and expressed as $IC_{50}$ values as described earlier. Four incubation times were employed for each inhibitor: 96 (standard assay time), 72, 48 and 24 h.

**Fluorescence microscopy**

*In vitro*, synchronised ring-stage parasite cultures were treated at 3x$IC_{50}$ for TAE684, AT83, and ZM-39 and at $IC_{99}$ for hesperadin and sampled every 12 hours over a 60-hour period. Parasites were then evaluated as follows:

**MitoTracker Viability:** For parasite viability, cells were then stained for 30 min with 150 nM MitoTracker Orange CMTMRos (M7510, ThermoFisher) and fixed using 4% paraformaldehyde (PFA), 0.075% glutaraldehyde (GA) for 15 min at 37 °C. Finally, cells were mounted on a glass slide using ProLong™ glass antifade mount medium with NucBlue™ (P36981, ThermoFisher). Images were captured using an EVOS M5000 (ThermoFisher).

**Immunofluorescence**: Nuclear content and microtubule morphology were assessed using direct immunofluorescence imaging on fixed parasites (4% PFA, 15 min) on poly-D-lysine-coated coverslips. These were washed three times with PBS, permeabilised with fresh 0.1% Triton X-100, washed another three times with PBS, and blocked with 3% BSA-PBS for 1 h at room temperature (RT). Coverslips were then exposed to 1:500 dilution primary anti-tubulin, mouse monoclonal (Merck, T5192) overnight at 4 °C. Primary antibody was subsequently washed off three times with PBS and incubated in anti-mouse conjugated CF488A secondary antibody produced in chicken (Merck, SAB4600238) for 1 h at RT. Cells were then mounted on a glass slide using ProLong™ glass antifade mount medium with NucBlue™ (P36981, ThermoFisher). Membrane morphology and nuclear material segregation were assessed using fixed parasites (4% PFA, 0.075% GA for 15 min) on poly-D-lysine-coated coverslips. Coverslips were washed three times with PBS and incubated overnight at RT with 5 µM BODIPY-TR-ceramide (D7540, ThermoFisher). Cells were then mounted on a glass slide using ProLong™ glass antifade mount medium with NucBlue™ (P36981, ThermoFisher).

**Image acquisition:** Images were acquired using a Zeiss LSM780 Inverted Confocal Laser Scanning Microscope (LSM) (Zeiss, Germany) for super-resolution imaging in the appropriate channels with a ×100 oil-immersion objective and 1.4 numerical aperture. Images were processed using Zeiss ZEN Lite Blue Edition software (Zeiss, Germany) and Fiji software.

**Expansion Microscopy:** For ultra expansion microscopy (U-ExM), the procedure was performed as previously described[10,11]. In short, tightly synchronised *Pf*NF54 cultures were treated with hesperadin (3 µM) and harvested at ± 46 hpi. Parasite cultures (0.5% hematorcrit) were then seeded onto and incubated for 30 min at 37 °C on 12 mm round Coverslips, treated with poly-D-lysine for 1 h at 37 °C, in the wells of a 12-well plate. Culture supernatants were removed, and cultures were fixed with 1 mL of 4% v/v PFA in 1× PBS for 15 min at 37 °C. Following fixation, coverslips were washed three times with 37 °C PBS before being treated with 1 mL of 1.4% v/v formaldehyde/2% v/v acrylamide (FA/AA) in PBS. Samples were then incubated at 37 °C overnight. Monomer solution (19% w/w sodium acrylate, 10% v/v acrylamide, 0.1% v/v N,N'-methylenebisacrylamide in PBS) was made the night before gelation and stored at −20 °C overnight. Prior to gelation, FA/AA solution was removed from coverslips, and they were washed once in PBS. For gelation, 5 µL of 10% v/v tetraethylenediamine (TEMED) and 5 µL of 10% w/v ammonium persulfate (APS) were added to 90 µL of monomer solution and briefly vortexed. Subsequently, 35 µL was pipetted onto parafilm, and coverslips were placed (cell side down) on top. Gels were incubated at 37 °C for 30 min before being transferred to wells of a 6-well plate containing denaturation buffer (200 mM SDS, 200 mM NaCl, 50 mM Tris, pH 9). Gels were incubated in denaturation buffer with shaking for 15 min, before the separated gels were transferred to 1.5 mL tubes containing denaturation buffer. 1.5 mL tubes were incubated at 95 °C for 90 min. Following denaturation, the gels were transferred to 10 cm Petri dishes containing 25 mL of MilliQ water for the first round of expansion and placed on a shaker for 30 min three times, with the water changed between each time. Gels were subsequently shrunk with two 15 min washes in 25 mL of 1× PBS, before being transferred to 6-well plates for 30 min of blocking in 3% BSA-PBS at room temperature. After blocking, the gels were incubated overnight with primary antibodies diluted in 3% BSA-PBS. After primary antibody incubation, gels were washed three times in 0.5% v/v PBS-Tween 20 for 10 min before incubation with secondary antibodies diluted in 1× PBS for 2.5 h. Following secondary antibody incubation, gels were again washed three times in PBS-Tween 20, before being transferred back to 10 cm Petri dishes for re-expansion with three 30 min MilliQ water incubations. Gels were either imaged immediately following re-expansion or stored in 0.2% w/v propyl gallate in MilliQ water until imaging.

**Image acquisition:** A small slice of gel (~10 × 10 mm) was cut and mounted on an imaging dish (35 mm Cellvis coverslip-bottomed dishes, Fisher Scientific) coated with poly-D-lysine. The side of the gel containing the sample is placed face down on the coverslip, and a few drops of MiliQ $H_2O$ are added after mounting to prevent gel shrinkage due to dehydration during imaging. All images presented in this study were captured using a Zeiss LSM980 microscope with an AxioObserver and an Airyscan 2 detector. Imaging was conducted on both microscopes using a ×63 Plan-Apochromat objective lens with a numerical aperture of 1.4. All images were acquired as Z-stacks that had an XY pixel size of 0.035 µm and a Z-slice size of 0.13 µm. Images were processed using Zeiss ZEN Lite Blue Edition software (Zeiss, Germany) and Fiji software.

**Microtubule branch length image processing:** Rolling ball background subtraction and Gaussian blur were first applied to the image channel to reduce noise. The microtubule branches were then semi-automatically traced using the FIJI Simple Neurite Tracer (SNT) software V4.2.1.[12].

**Inhibitor susceptibility assays using *P. falciparum* kinase cKD lines**

Compound susceptibility assays using *P. falciparum* Ark1 and Ark2 cKD lines were carried out as previously described[13]. Ark1 (PF3D7_0605300) and Ark2 (PF3D7_0309200) cKD lines generated by fusing the coding sequence and non-coding RNA aptamer sequences in the 5'- and 3'-UTR, permitting translation regulation using the TetR-DOZI system [14]. Editing was achieved by CRISPR/SpCas9 using the linear pSN054 vector that contains cloning sites for the left homology region (LHR) and the right homology region (RHR), as well as a gene-specific guide RNA under control of the T7 promoter[15]. The final constructs were sequence-verified and further confirmed by restriction digests. Transfection into Cas9- and T7 RNA polymerase-expressing NF54 parasites was carried out by pre-loading erythrocytes with the donor vector as previously described[16]. Parasite culture was maintained continuously in 500 nM anhydrotetracycline (aTc, Sigma-Aldrich 37919) and drug selection with 2.5 µg/mL of Blasticidin S (RPI Corp B12150-0.1) was initiated four days after transfection.

Briefly, synchronous ring-stage Ark1 (PF3D7_0605300) and Ark2 (PF3D7_0309200) cKD parasites, as well as a control parasite line expressing an aptamer-regulatable fluorescent protein, were maintained in the presence of high aTc (500 nM) or no aTc and distributed into 384-well polystyrene microplates (Corning). Stock solutions of compounds were serially diluted and transferred to the parasite-containing plates using the Janus platform (PerkinElmer). DMSO and dihydroartemisinin treatment (500 nM) served as reference controls. Luminescence was measured after 72 hours using the Renilla-Glo Luciferase Assay System (Promega E2750) and the GloMax Discover Multimode Microplate Reader (Promega), and $IC_{50}$ values were obtained from corrected dose-response curves using Graph-Pad Prism.

***In vitro Plasmodium* kinase inhibition assays:**

***Pf*Ark1/3 (Luceome Biotechnologies)**

A 10 mM stock of the inhibitor was diluted in DMSO to a concentration of 25 µM. Prior to initiating a profiling campaign, the inhibitors were evaluated for false positive against split-luciferase. The inhibitors were then screened in duplicate against each of the kinases. For kinase assays, each Cfluc-Kinase was translated alongside Fos-Nfluc using a cell-free system (rabbit reticulocyte lysate) at 30 °C for 90 min. A 24 µL aliquot of this lysate containing either 1 µL of DMSO (for the no-inhibitor control) or the inhibitor solution in DMSO (at a final concentration of 1 µM) was incubated for 2 hours at room temperature in the presence of a kinase-specific probe. An 80 µL volume of luciferin assay reagent was added to each solution, and luminescence was immediately measured using a luminometer. The% Inhibition and% Activity Remaining were calculated using the following equations:% Inhibition = (ALUControl – ALUSample) x 100 / ALUControl and% Activity Remaining = 100 -% Inhibition. Profiling data for all kinases were plotted as% activity remaining versus the kinases profiled. A heat map representing the effect of the inhibitor on the kinases was also generated.

***Pf*PKG**

Full length *Pf*PKG (PF3D7_1436600) was expressed in *E. coli* Rosetta 2 (Novagen, EMD_BIO-71402) as previously described [17–19]. Briefly, the N-terminal His-tagged recombinant *Pf*PKG protein was purified using a HisTrap HP column (GE Healthcare), followed by anion exchange and size exclusion chromatography (HiLoad 16/600 Superdex 200 pg column, GE Healthcare). Final buffer composition of purified protein was 50 mM Tris-HCl pH 8.0, 150 mM NaCl, 10 mM β-mercaptoethanol, 10% glycerol. *Pf*PKG $IC_{50}$ assays were performed based on previously described methods using the ADP-Glo Kinase Assay (Promega) to measure ADP formation. Briefly, a 3-fold serial dilution of each inhibitor was carried out in DMSO and inhibitors were subsequently diluted into assay buffer (25 mM HEPES pH 7.4, 0.1 mg/mL BSA, 0.01% (v/v) Triton-X 100, 20 mM $MgCl_2$, 2 mM DTT, 10 µM cGMP) to 1.5 × the final required concentration. 2 µL of each inhibitor dilution was transferred into a white 384-shallow well plate (Nunc #264706). A MANTIS® Liquid Handler (Formulatrix) was used to dispense the remaining assay components. 0.5 µL *Pf*PKG protein was added and following a 5-min pre-incubation with inhibitor, 0.5 µL substrate buffer (ATP and peptide substrate GRTGRRNSI-$NH_2$), was added to each well. The final 3 µL kinase reaction contained ~1 nM *Pf*PKG protein, 10 µM ATP, 20 µM GRTGRRNSI-$NH_2$, 1% (v/v) DMSO and inhibitor in assay buffer. Reactions were incubated for 45 min at 22 °C (resulting in < 10% ATP conversion). ADP formation was measured using the ADP-Glo Kinase Kit (Promega). Briefly, 2 µL ADP-Glo reagent was added to each well and incubated for 40 min at 22 °C to deplete the remaining ATP. 2 µL of Kinase Detection Reagent was then added, and the reaction was incubated for a further 30 min at 22 °C. The plate was sealed with an adhesive foil seal for all incubation steps.

Luminescent signal was measured using the EnSpire Multimode Plate Reader (PerkinElmer). The data was normalised based on the 100% activity controls (1% DMSO only) and the 100% inhibition controls (10 µM ML10 (N-[5-[3-[2-(cyclopropylmethylamino)pyrimidin-4-yl]-7-[(dimethylamino)methyl]-6-methylimidazo[1,2-a]pyridin-2-yl]-2-fluorophenyl]methanesulfonamide), LifeArc). Mean $IC_{50}$ values were calculated from N = 2 independent experiments, each with technical duplicates (log(inhibitor) vs. normalised response - Variable slope). $IC_{50}$ values within 3-fold of independent experiments are considered reproducible.

***Pf*CLK3**

The kinase domain of *Pf*CLK3 (PF3D7_1114700, Gln317 to Ser694) was expressed in *E. coli* Rosetta™(DE3)pLysS cells (Novagen, EMD_BIO-70956). The expression construct and protein production protocol were kindly provided by Rafael M. Couñago, CQMED, University of Campinas. Briefly, the N-terminal His-tagged recombinant PfCLK3 protein was purified using a HisTrap HP column (GE Healthcare), and size exclusion chromatography (HiLoad 16/600 Superdex 200 pg column, GE Healthcare). Final buffer composition of purified protein was 20 mM HEPES, pH 7.5, 0.5 M NaCl, 5% glycerol and, 10 mM β-mercaptoethanol. *Pf*CLK3 $IC_{50}$ assays were performed using the ADP-Glo Kinase Assay (Promega) to measure ADP formation. Briefly, a 3-fold serial dilution of each inhibitor was carried out in DMSO, and inhibitors were subsequently diluted into assay buffer (50 mM HEPES pH 7.4, 0.1 mg/mL BSA, 0.01% (v/v) Triton-X 100, 20 mM $MgCl_2$, 2 mM DTT) to 1.5 × the final required concentration. 2 µL of each inhibitor dilution was transferred into a white 384-shallow well plate (Nunc #264706). A MANTIS® Liquid Handler (Formulatrix) was used to dispense the remaining assay components. 0.5 µL *Pf*CLK3 protein was added and following a 30-minute incubation with inhibitor, 0.5 µL substrate buffer (ATP and peptide substrate Myelin Basic Protein (MBP), was added to each well. The final 3 µL kinase reaction contained ~15 nM *Pf*CLK3 protein, 10 µM ATP, 20 µM MBP, 1% (v/v) DMSO and inhibitor in assay buffer. Reactions were incubated for 75 min at 30 °C (resulting in < 10% ATP conversion). ADP formation was measured using the ADP-Glo Kinase Kit (Promega) as described for *Pf*PKG. The data was normalised based on the 100% activity controls (1% DMSO only) and the 100% inhibition controls (10 µM TCMDC-135051[20]), and data were analysed as described for *Pf*PKG.

***Pv*PI4Kβ**

Full-length *Pv*PI4Kβ (PVX_098050) recombinant protein was expressed in a baculovirus-insect cell expression system and purified as previously described[17,21]. Briefly, N-terminal His-tagged recombinant *Pv*PI4Kβ protein was purified using a HisTrap HP column (GE Healthcare), followed by size exclusion chromatography (HiLoad 16/600 Superdex 200 pg column, GE Healthcare). Final buffer composition of purified protein was 20 mM HEPES pH 7.5, 500 mM NaCl, 5% (v/v) glycerol, 10 mM β-mercaptoethanol. *Pv*PI4Kβ kinase inhibition assays were performed using the ADP-Glo kinase assay kit (Promega) to measure ADP formation. L-alpha-phosphatidylinositol (PI; Avanti Polar Lipid, cat. 840042P) dissolved in 3% n-Octylglucoside to a stock concentration of 20 mg/mL was used as the lipid substrate. Briefly, a 3-fold serial dilution of each inhibitor was carried out in DMSO, and inhibitors were subsequently diluted into assay buffer (25 mM HEPES pH 7.4, 100 mM NaCl, 3 mM $MgCl_2$, 1 mM DTT, 0.025 mg/ml BSA, 0.2% (v/v) Triton-X-100) to 1.5 × the final required concentration. 2 µL of each inhibitor dilution was transferred into a white 384-shallow well plate (Nunc #264706). A MANTIS® Liquid Handler (Formulatrix) was used to dispense the remaining assay components. 0.5 µL *Pv*PI4Kβ protein was added and following a 5-min pre-incubation with inhibitor, 0.5 µL substrate buffer (ATP and PI) was added to each well. The final 3 µL kinase reaction contains ~6 nM *Pv*PI4Kβ protein, 10 µM ATP, 0.1 mg/ml PI, 1% (v/v) DMSO and inhibitor in assay buffer. Reactions were incubated for 40 min at 22 °C (resulting in < 10% ATP conversion). ADP formation was measured using the ADP-Glo Kinase Kit (Promega) as described for *Pf*PKG, the only difference being that 10 mM $MgCl_2$ was added to the ADP-Glo reagent prior to use. *The Pv*PI4Kβ inhibitor MLN0128 (sapanisertib[22]) at 10 µM was used as the control (100% inhibition), and data were analysed as described for *Pf*PKG.

**Molecular docking.**

A homology model for *P. falciparum* Ark1 (*Pf*Ark1) was generated using the crystal structure of human Aurora kinase A (*Hs*AurA) co-crystallized with ATP (5DNR)[23], with a 33% sequence identity. The protein sequences of the *Pf*Ark1, *Hs*AurA, *Hs*AurB and *X. laevis* AurB (*Xen*AurB) were aligned using the NCBI alignment tool[24]. Ligands were prepared on the Schrödinger 2021-4 Maestro interface using the LigPrep tool at pH 7.4 ± 0.5 using the OPLS4 forcefield, while the crystal was prepared using the Schrödinger Protein Preparation Wizard. Structural optimization included bond order assignments, generation of tautomer and ionization states, addition of hydrogen atoms, side chains and loops, removal of water molecules from the surface of the crystal structure and restrained minimization at pH 7.4 with the default parameters to generate the lowest energy conformation using the OPLS4 forcefield[25,26]. The receptor grid was generated for the refined *Pf*Ark1 homology model structures by centering a 10 Å inner grid box and a 20 Å outer grid box over specified key residues found in the hinge region (Phe107-The119). Ligands were docked to the receptor using the Glide default settings in extra precision, flexible ligand mode, and poses were scored using the Schrödinger GlideScore function[27]. Following this, further analysis of the docking poses was conducted using Molecular Mechanics and Generalized Born Surface Area (MM-GBSA) with the default solvation model (VSGB and OPLS4) settings, and residues within 20 Å of the active site were allowed to be flexible[28]. Docking poses were visualized and images were generated in Schrödinger 2021-4 Maestro interface.

**Isobologram analysis**

Fixed-ratio isobologram analysis against asexual intra-erythrocytic parasites was performed as previously described in[29]. Parasites were treated with fixed ratios (5:0, 4:1, 3:2, 2:3, 1:4, and 0:5) for TAE684 and hesperadin in combination with chloroquine (CQ), ML10, and each other similarly, this was done for AT83 and ZM-39 in combination with CQ and with one another using the SYBR Green I proliferative assay[4] as readouts to determine the fractional inhibitory concentrations (FIC) for the respective combinations. Isobolograms were generated by plotting paired FIC values linearly, utilising the average of three biological replicates, each performed in technical duplicates. The paired FIC values for each drug combination were analysed, and the mean FIC values (ΣFIC) were calculated to delineate synergism (<0.8), additivity/indifference (0.8-1.4), or antagonism (>1.4).

**NP-40-mediated cell-free β-hematin inhibition assay:**

The β-hematin inhibition assay, as described in[30,31] was used to test the inhibitors for their ability to inhibit β-hematin formation. The test samples were prepared as 20 mM or 10 mM stock solutions in dimethyl sulfoxide (DMSO). Serial dilutions of each inhibitor (100 µL) were performed from column 12 to column 2 of a 96-well plate in triplicate, with column 1 serving as a negative control (0 µM test inhibitor). The samples were tested at a starting in-well concentration of 1000 µM (20 mM stock) or 500 µM (10 mM stock). An "NP-40 substitute" detergent was added (305.5 µM) to each well to mimic the lipophilic environment in which hemozoin formation occurs within the parasite's digestive vacuole, in helping to mediate the formation of β-hematin. A 25 mM hematin stock was prepared by sonicating heme in DMSO and then suspending 178.8 µL of this in 1 M acetate buffer (20 mL, pH 4.8). Then, 100 µL of the hematin suspension was added to each well to a final well volume of 200 µL. The plate was then incubated for 5 hours at 37 °C. After incubation, 32 µL and 60 µL of 50% pyridine solution and acetone were added, respectively. The pyridine-ferrochrome method by[31] was used for UV-vis analysis, with the heme-pyridine complex absorbance measured at 405 nm. Data were analysed using Microsoft Excel and GraphPad Prism v10.5 software.

**Heme fractionation assay:**

The assay set-up followed that described in[30,32] optimised to a multi-well colourimetric assay for determining heme species in *P. falciparum*. Briefly, *P. falciparum* ring-staged parasites were incubated at 2x $IC_{50}$ value of the test inhibitor for 24 hours. Thereafter, the incubated parasites (trophozoites) were harvested and isolated through various steps of a cellular fractionation process. The cellular fractionations allow for direct quantification of the three major heme species in the trophozoites, namely hemoglobin, free heme, and hemozoin. These heme species can be determined spectroscopically using the aqueous pyridine-ferrochrome method. This method is based on the principle that aqueous pyridine forms a low-spin complex with heme but not hemozoin, and since the absorbance obeys Beer's law, it allows for the quantification of heme concentration in solution. The various heme fractions were recovered and measured as follows, after the *P. falciparum* trophozoite cells were harvested following exposure to probe inhibitors:

**Hemoglobin fraction:**

Water (100 µL) was added to lyse the cells, and the suspension was sonicated for 5 min. This was followed by the addition of 50 µL 0.2M HEPES buffer pH 7.5, and then centrifugation at 3600 rpm for 20 min. The resulting supernatant (containing hemoglobin) was then transferred into an adjacent set of wells on the same plate. To which 50 µL of each of 4% SDS, 0.3 M NaCl and 25% pyridine were added, respectively. Thereafter, 200 µL of the resulting solution (400 µL) was transferred to a set of wells on a separate 96-well plate for UV-vis analysis[32,33].

**Heme fraction:**

The pellet from the centrifugation step described above consists of heme and hemozoin. Notably, it is known that pyridine dissolves heme to form a low-spin complex, while the hemozoin remains and can be spun out. Hence, to the pellet, 50 µL of water and 50 µL 4% SDS was added, resuspended and sonicated for 5 min. Then 50 µL each of HEPES 0.2 M pH 7.5, 0.3 M NaCl, and 25% pyridine were added, respectively. The mixture was centrifugated at 3600 rpm and the supernatant (250 µL) was transferred to adjacent wells on a separate 96-well plate. Thereafter, 150 µL of water was added and 200 µL of the mixture transferred to a set of wells on a separate plate for UV-vis analysis[32,33].

**Hemozoin fraction:**

To the remaining pellet, 50 µL each of water and 0.3 M NaOH were added, and the mixture was sonicated for 15 min. After 30 min of incubation at room temperature, 50 µL each of HEPES 0.2 M pH 7.5, 0.3 M HCl, and 25% pyridine were added, respectively. To the resulting mixture (250 µL), 150 µL of water was then added, and 200 µL of this was transferred to a set of wells on a separate 96-well plate for UV-vis analysis[32,33].

**Cell counting using flow cytometry**

Cell counts for the assay were performed using the flow cytometry method described in[33]. Cell counting determines the number of trophozoites per sample and, therefore, allows for the quantification of the amount of heme found in each individual cell.

**Statistical analysis**

A two-tailed t-test (95% CI) was used for the statistical assessment of measurement differences that might present significance relative to the control. The significant differences are displayed as asterisks on graphs: * $p < 0.05$; ** $p < 0.01$; *** $p < 0.001$, ****$p < 0.0001$. The data presented was from one experiment performed in quadruplicate and subsequently analysed using Microsoft Excel and GraphPad Prism v10.5 software.

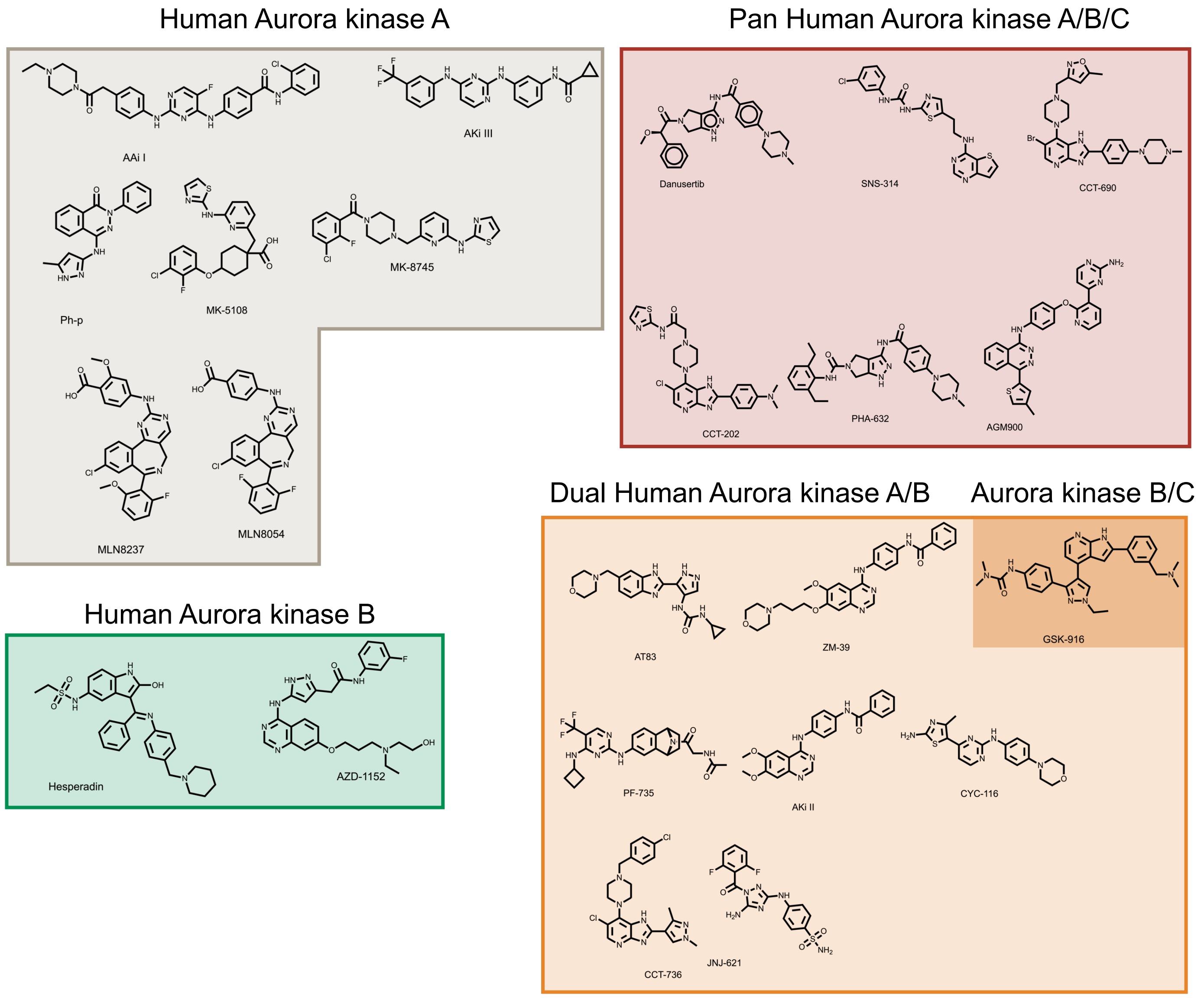


**Supplementary Figure S1:**
Structures of the selected Aur class anticancer inhibitors, classified according to their human aurora kinase inhibitor class.

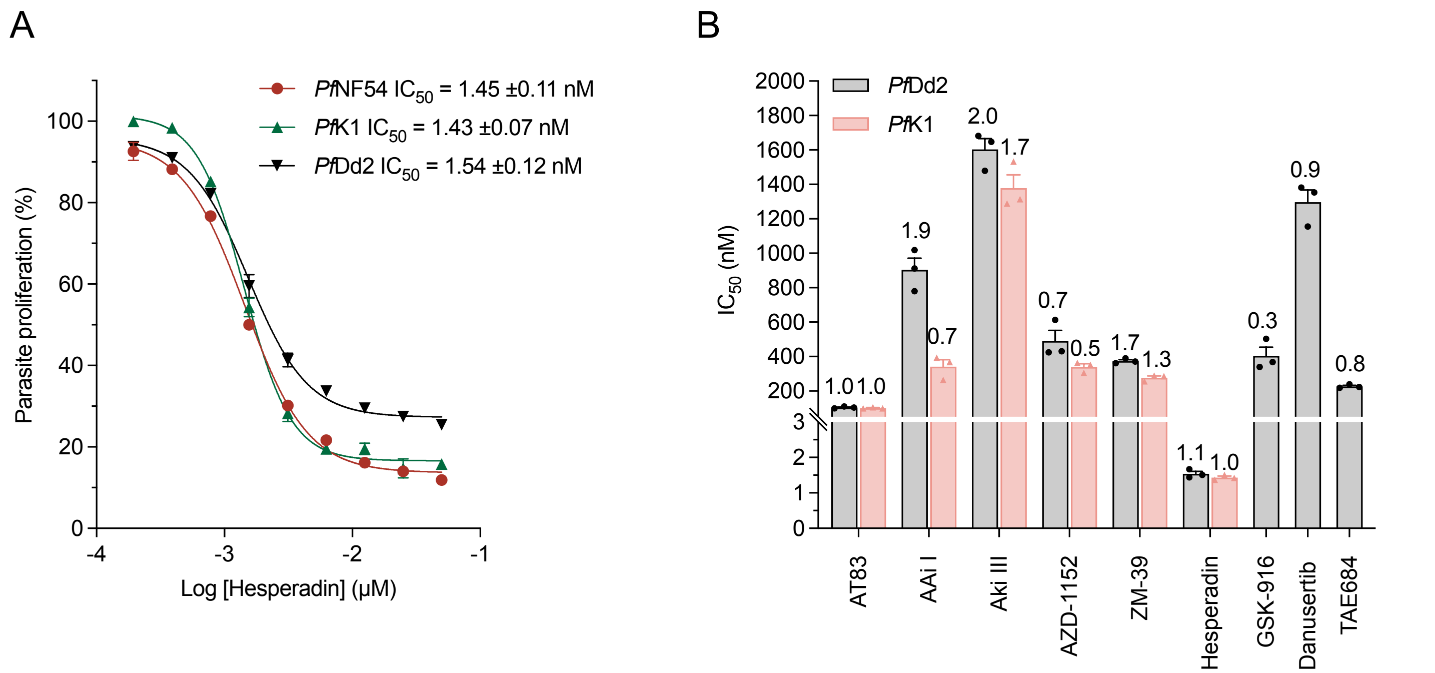


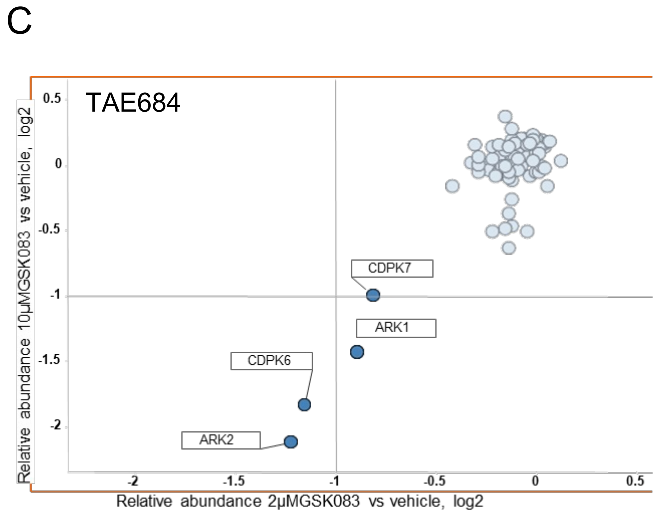


| PlasmoDB code | Protei description | Quant. Uniq. Peptides | Folde change compared to vechicle control (+10 μM GSK2122083A) |
|---|---|---|---|
| PF3D7_0309200 | serine/threonine protein kinase, ARK2 | 14 | 0.23 |
| PF3D7_1122800 | calcium dependent protein kinase 6 | 9 | 0.28 |
| PF3D7_0605300 | serine/threonine protein kinase, ARK1 | 10 | 0.37 |
| PF3D7_1123100 | calcium-dependent protein kinase 7 | 23 | 0.5 |

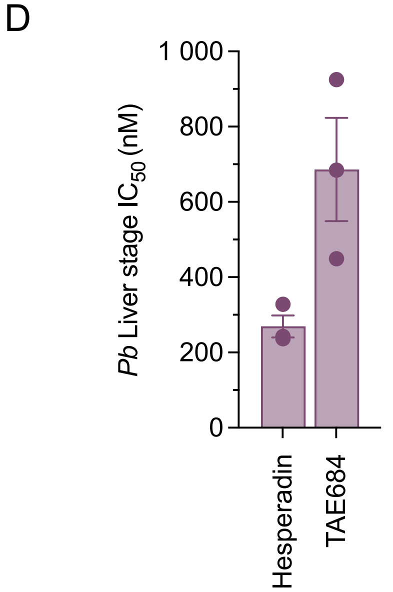


**Supplementary Figure S2:**
**A)** Dose-response curve of Hesperadin against ABS *Pf*NF54, *Pf*Dd2 and *Pf*K1. **B)** Antiplasmodial activity of nine anticancer aurora kinase inhibitors (selected based on their *Pf*NF54 ABS activity) against *P. falciparum* multidrug-resistant ABS parasites (*Pf*Dd2 and *Pf*K1). **C)** Kinobead competitive inhibition data. A total of three *Plasmodium* kinases were competed from Kinobeads by TAE684 at 10 µM with a $log_2$ fold change below the cutoff of -1 relative to the vehicle control. **D)** Prophylactic antiplasmodial activity ($IC_{50}$) of Hesperadin, TAE684 and Danusertib in an *in vitro P. berghei* ANKA liver stage assay (n=3, mean ± S.E.).

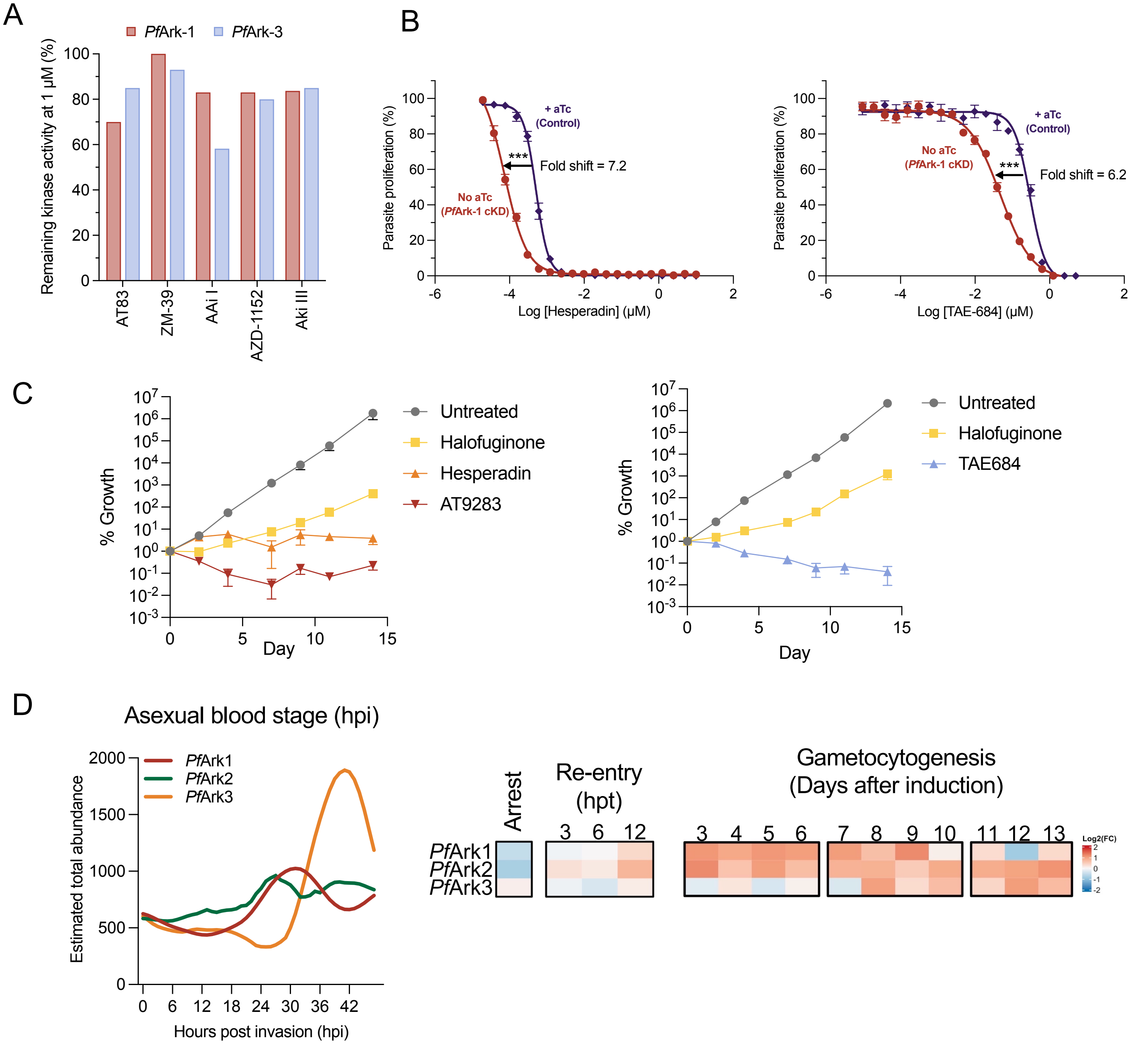


**Supplementary Figure S3:**

**A)** Single-point activity profile of selected inhibitors at 1 μM against recombinant *Pf*Ark-1 and *Pf*Ark-3 proteins, utilising a three-hybrid split-luciferase competitive binding assay (KinaseSeeker™). **B)** Effect of conditional knockdown (cKD) of *Pf*Ark-1 on parasite sensitivity to Hesperadin and TAE-684, relative to control conditions in the presence of high aTc. Representative dose-response curves are presented for each cKD parasite line ($n$ = 3, mean ± S.E.) with an unpaired two-tailed t-test. **C**) Cumulative growth profiles for drug-treated and no-drug controls. **D)** Expression profile of *Pf*Ark members across ABS, gametocytogenesis, as well as during G1/S-like arrest.

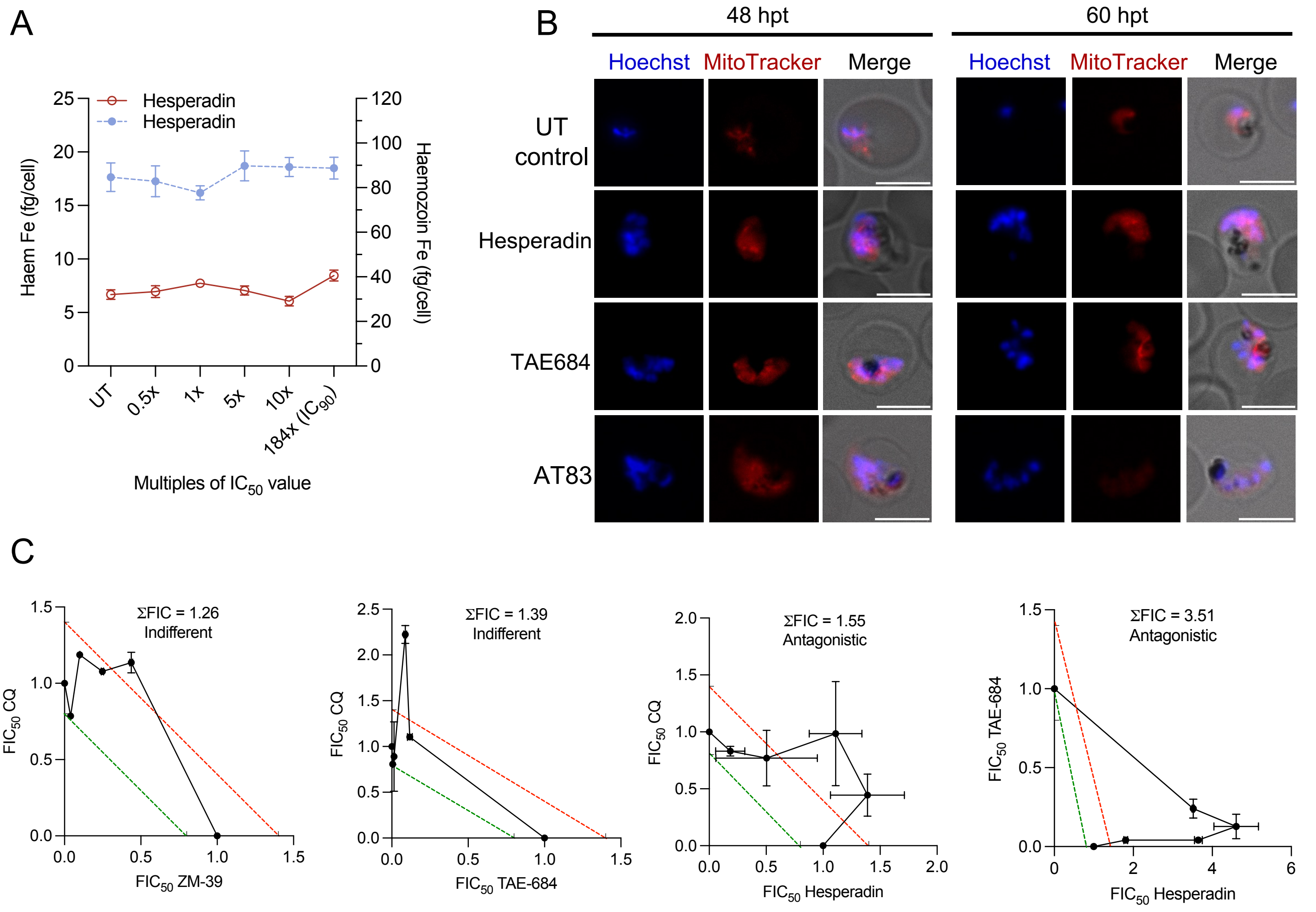


**Supplementary Figure S4:**

**A)** Dose-dependent changes in the heme Fe levels from intracellularly-extracted fractions of hemozoin under hesperadin treatment. **B)** MitoTracker viability of treated parasites 48 and 60 hpt. Scale bar = 5 μm. **C)** Fixed-ratio isobologram analysis for selected inhibitors in various combinations against asexual intra-erythrocytic parasites. Data represent the mean fractional inhibitory concentration ($FIC_{50}$) of two biological replicates; each performed in technical duplicates. Error bars represent ± S.E.

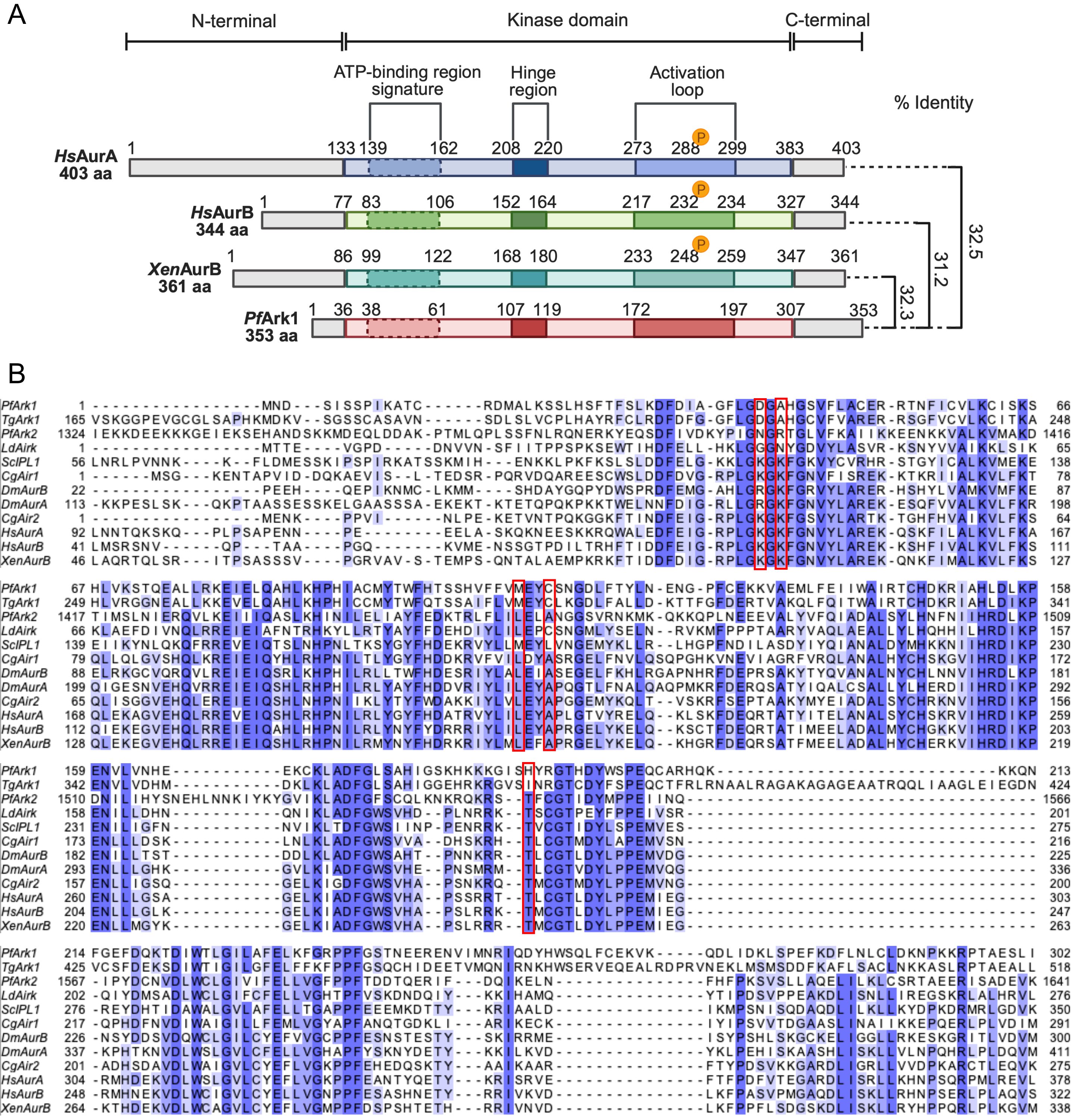


**Supplementary Figure S5:**

**A)** Schematic representation of *P. falciparum* Ark1 and -2 and sequence alignment of conserved motifs of mammalian aurora kinases. **B)** Sequence alignment of other eukaryotic organisms and *P. falciparum* aurora kinase proteins. Identity/similarity is indicated with the blue shading, and red boxes indicate critical changes in the *Plasmodium* protein.

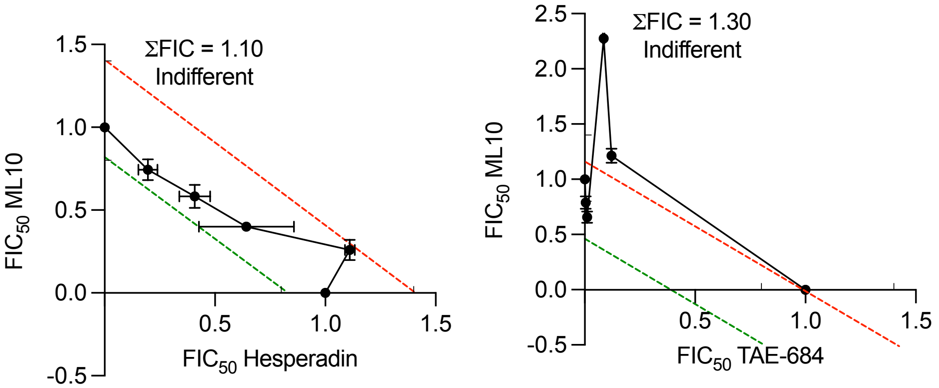


**Supplementary Figure S6:**

Fixed-ratio isobologram analysis for selected inhibitors in various combinations against asexual intra-erythrocytic parasites. Data represent the mean fractional inhibitory concentration ($FIC_{50}$) of two biological replicates; each performed in technical duplicates. Error bars represent ± S.E.

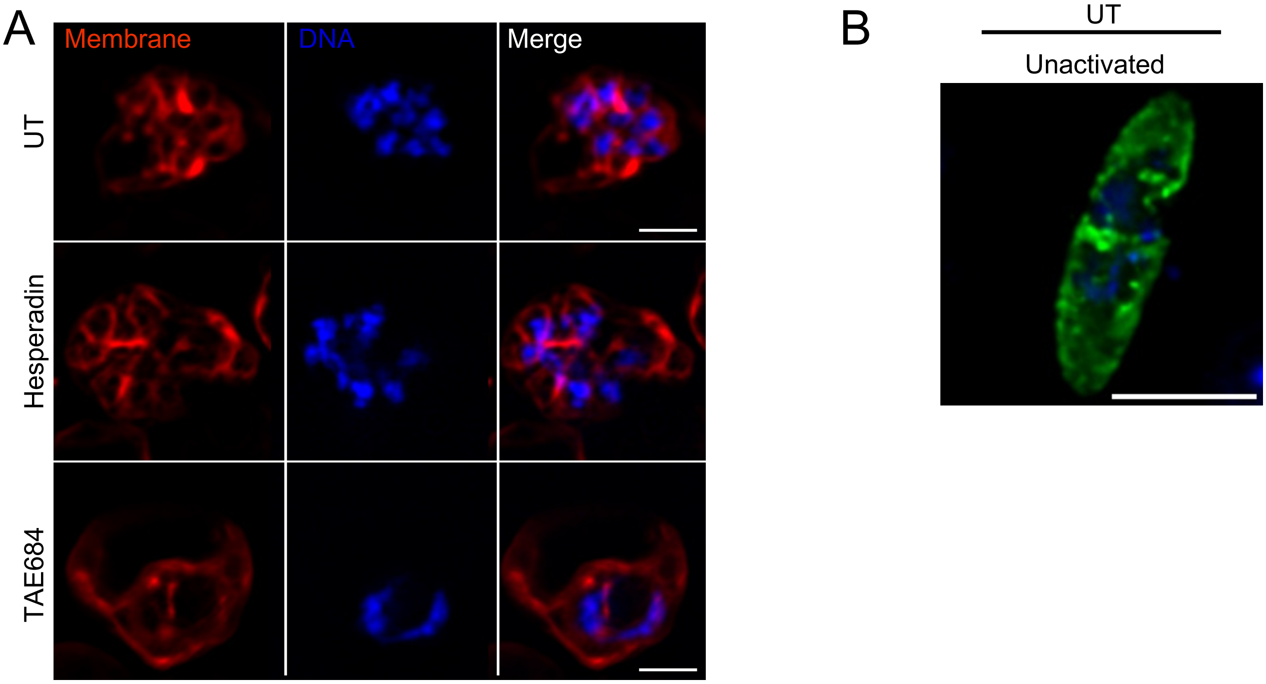


**Supplementary Figure S7:**
**A)** Representative images (average intensity projections) of the morphological effect observed on membrane formation in hesperadin and TAE684-treated samples. The images represent at least ten parasites per sample. Scale bars correspond to 2 µm. **B)** A representative image of a non-activated mature gametocyte. Scale bar = 5 µm.
